# AlphaDIA enables End-to-End Transfer Learning for Feature-Free Proteomics

**DOI:** 10.1101/2024.05.28.596182

**Authors:** Georg Wallmann, Patricia Skowronek, Vincenth Brennsteiner, Mikhail Lebedev, Marvin Thielert, Sophia Steigerwald, Mohamed Kotb, Tim Heymann, Xie-Xuan Zhou, Magnus Schwörer, Maximilian T. Strauss, Constantin Ammar, Sander Willems, Wen-Feng Zeng, Matthias Mann

**Affiliations:** Proteomics and Signal Transduction, Max Planck Institute of Biochemistry, Martinsried, Germany; Proteomics Program, Novo Nordisk Foundation Center for Protein Research, Faculty of Health and Medical Sciences, University of Copenhagen, Copenhagen, Denmark

## Abstract

Mass spectrometry (MS)-based proteomics continues to evolve rapidly, opening more and more application areas. The scale of data generated on novel instrumentation and acquisition strategies pose a challenge to bioinformatic analysis. Search engines need to make optimal use of the data for biological discoveries while remaining statistically rigorous, transparent and performant. Here we present alphaDIA, a modular open-source search framework for data independent acquisition (DIA) proteomics. We developed a feature-free identification algorithm particularly suited for detecting patterns in data produced by sensitive time-of-flight instruments. It naturally adapts to novel, more eTicient scan modes that are not yet accessible to previous algorithms. Rigorous benchmarking demonstrates competitive identification and quantification performance. While supporting empirical spectral libraries, we propose a new search strategy named end-to-end transfer learning using fully predicted libraries. This entails continuously optimizing a deep neural network for predicting machine and experiment specific properties, enabling the generic DIA analysis of any post-translational modification (PTM). AlphaDIA provides a high performance and accessible framework running locally or in the cloud, opening DIA analysis to the community.

## Introduction

Proteomics entails the study of key players of life – proteins – and their translation, composition of isoforms, post-translational modification and degradation^1^. As proteomes are composed of thousands of different proteoforms, which produce hundreds of thousands of peptides in bottom-up proteomics, handling complexity is central to MS based proteomics acquisition and bioinformatic analysis.

Until recently, data dependent acquisition (DDA) was the acquisition method of choice. The direct relationship between selected precursors and relatively pure fragmentation spectra, combined with its mature ecosystem of search engines, results in confident peptide identifications^2–5^. Due to the straightforward relationship between precursor and fragment spectrum, this also holds for challenging cases such as complex patterns of post-translational modifications or the interpretation inter-protein cross-links^6,7^. Yet, selecting only a single peptide at a time comes at the cost of increased data acquisition time and stochastic sampling of precursors across liquid chromatography (LC)-MS runs^8^.

In contrast to DDA, Data Independent Acquisition (DIA), allows the selection of multiple peptides in parallel, originally in the form of cycles of fixed-width, relatively wide selection windows^9,10^. This results in systematic sequencing of all available peptides only limited by sensitivity. Importantly, repeated scanning of the same mass range yields complete elution profiles of both the precursors and the fragments. This increases dynamic range, allows for faster acquisition and deeper proteome characterization down to the single cell level^11,12^. The principal challenge of DIA is the increased spectral complexity as multiple peptides fragment together leading to convoluted spectra. Thus, DIA data by its nature requires algorithms to deconvolute overlapping fragmentation patterns and assign peptide identifications.

Initially, DIA involved generating an empirical, sample specific spectral library, usually acquired by offline fractionation of samples and DDA acquisition, or spectrum centric processing^13,14^. Different algorithms have been designed to process DIA data. Deconvolution of co-isolated peptides into individual spectra effectively reduces them to DDA like data, amenable to the plethora of proven DDA methods. However, peptide-centric approaches, in which each spectrum of the library is matched to the complex DIA data, achieve higher performance especially if paired with deep-learning based scoring of identifications as pioneered by Demichev et al. ^15–17^. Deep learning also allows the prediction of libraries in silico, obviating the need for sample specific empirical libraries ^18–20^. However, for optimal performance this has so far required DDA data on the same MS platform and experimental method. This is in particular the case for spectra of post-translationally modified peptides^21,22^.

Despite the enormous potential of DIA, the fact that spectra are not easily manually interpretable has hindered full acceptance, especially as researchers must generally rely on few closed source algorithms. Flexible and open algorithms would clearly be beneficial to extend the reach, transparency, and acceptance of DIA. This becomes especially necessary as the most recent generation of instrument employs time-of-flight (TOF) detectors which are sensitive down to the single molecule level^23,24^. Raw files easily contain billions of detector events, often with no clearly visible peaks and up to four dimensions (4D) of separation^25^. Handling this data has usually required data reduction such as centroiding of the ion mobility, introducing feature boundaries or centroiding^26,27^, which may all lead to loss of information. We have found that this presents formidable challenges when implementing novel scan modes that make data processing even more demanding^28^, especially when the underlying algorithms and source code are not available.

To enable open, performant, and extensible processing of high complexity DIA data, we therefore propose a new processing framework which builds on technology driving the current breakthroughs in artificial intelligence, especially deep learning. Our algorithms view a DIA experiment as high-dimensional snapshot of the peptide spectrum space. This representation is amenable to DIA methods on all major instrument platforms and naturally covers simple DIA methods as well as ion mobility, variable windows, sliding quadrupole windows and yet to be developed acquisition modes. Integral to this generalized representation, the data is processed without reduction of retention time or mobility resolution. Instead, our feature-free approach performs machine learning directly on the raw signal, combing all available information before making discrete identifications. Furthermore, we propose an end-to-end deep transfer learning strategy based on our recently published alphaPeptDeep library. Transfer learning adapts the peptide library directly to the instrument and sample workflow. We showcase performance and versatility by extending DIA arbitrary PTMs, closing the gap between the versatility of DDA and the performance of DIA

## Results

We present alphaDIA, a modular, open-source, next generation framework for DIA search. It builds on the scientific python stack and the alphaX^29^ ecosystem allowing flexible search strategies as well as default workflows accessible through a Python API, Jupyter notebooks, a command line interface or an easily installable graphical user interface (**Fig. 1, a, Methods**). AlphaDIA covers the entire workflow from raw files to reporting protein quantities and can process files and proprietary formats from all major vendors. It was designed for ‘one stop processing’ of large cohorts and arbitrary data sizes, running natively on Windows, Linux and Mac or in a distributed fashion in the cloud with Slurm or Docker.

**Fig. 1:**
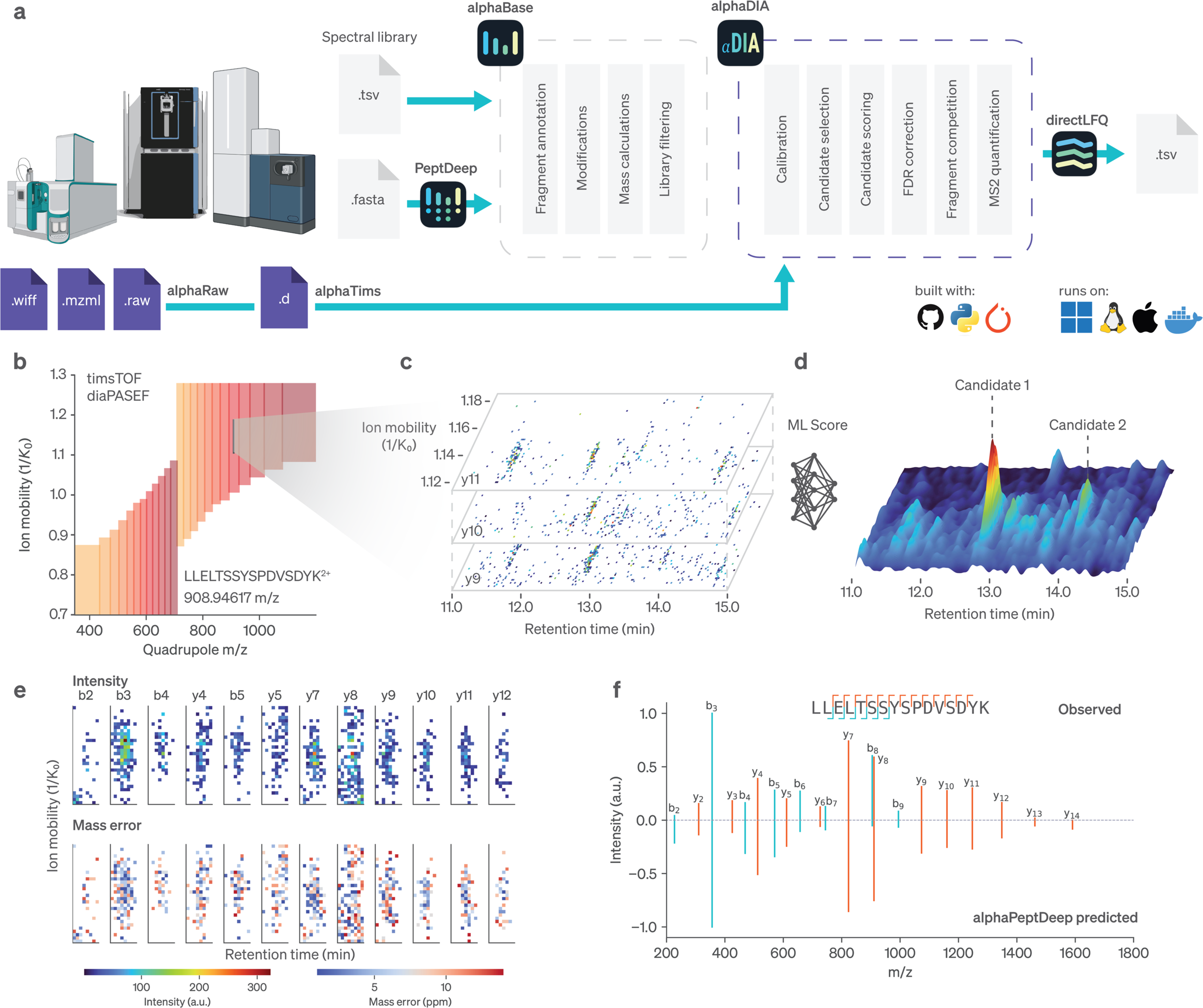
Overview of the alphaDIA framework. **a**, Components of alphaDIA and the integration into the alphaX ecosystem. AlphaDIA uses alphaRaw and alphaTims^30^ for accessing raw data from all major vendors. Importing as well as prediction of spectral libraries is facilitated by alphaBase and alphaPeptDeep^20^. After successful search, label free quantification is performed using directLFQ^31^. AlphaDIA uses best software engineering practices and builds on modern open architectures (GitHub, Python, PyTorch) **b-f**, TIMS DIA data acquired using optimal dia-PASEF^32^ is searched using a peptide centric algorithm. **b**, The library entry for a single peptide sequence is selected for search **c**, Fragment spectra containing the precursor of interest are extracted and converted into a dense matrix in spectrum space. **d**, Information from fragments mapping to the precursor of interest are combined in a continuous score. **e**, AlphaDIA defines candidate peak groups with discrete integration boundaries (top row: intensities, bottom row: mass deviation from theoretical mass. **f**, Aggregating signal across the integration boundaries in ion mobility and retention time reveals the peptide spectrum. For further scoring, AlphaPeptDeep spectrum predictions are used.

### Feature-free processing for high dimensional TOF data

Apart from state-of-the-art DIA processing, the impetus for alphaDIA was the shift towards fast, sensitive but also stochastic TOF detectors, presenting novel algorithmic challenges and opportunities. AlphaDIA’s feature-free and peptide-centric search is illustrated by the identification of the peptide LLELTSSYSPDVSDYK^2+^ from timsTOF Ultra dia-PASEF data (**Extended Data Fig. 1**). First, we select all MS1 and MS2 spectra that contribute evidence for this precursor (**Fig. 1,b**). A dense representation of the spectrum space is used to score potential peak group candidates, which does not involve feature building or centroiding (**Fig. 1,c-d**). Instead, signals are aggregated across retention time, ion mobility and fragments using learned convolution kernels. Only after all this evidence has been collected, discrete peak groups are determined (**Fig. 1, e**). In this way noisy TOF data in which individual fragment signals are not distinguishable from background can still be processed (**Extended Data Fig. 2**). After the signals in the peak groups are integrated it becomes evident that they correspond to a confidently identified peptide, given the agreement with the predicted spectrum (**Fig. 1,f**).

### Deep learning based search allows for whole proteome characterization

AlphaDIA uses deep learning based target-decoy competition and iterative calibration to search complex proteomes with spectral libraries. For each target precursor entry with a given sequence and charge state, a paired decoy peptide is created using a mutation pattern (**Methods**). Each peak group is scored by a collection of up to 47 features using a fully connected neural network (NN) (**Fig. 2, a**). False precursor identifications are controlled using a count-based FDR, calculated from the probabilities predicted by the NN (**Fig. 2, b-c**). Measured properties like retention time, ion mobility and m/z ratios are iteratively calibrated to the observed data on a high confidence subset of precursors, using non-linear LOESS regression with polynomial basis functions (**Fig. 2, d-f, Extended Data Fig. 3**). AlphaDIA uses spectrum centric fragment competition to ensure that fragment information is only used for a single precursor identification, even when multiple library entries match the same observed signal (**Methods**). On a 21 minute, 60 samples per day (SPD) gradient of HeLa cell lysate measured on a timsTOF Ultra with dia-PASEF, our algorithm identified more than 73,000 precursors with unique sequence and charge, corresponding to almost 6,800 protein groups (**Fig. 2, g-i**). For label free quantification (LFQ) we integrated the recently developed directLFQ algorithm^31^, which resulted in a median coefficient of variation of 7.7% for protein groups and a Person R > 0.99 across replicates (**Fig. 2, j-k**). This suggests that alphaDIA can search and quantify complex protein mixtures with excellent depth and quantitative precision.

**Fig. 2:**
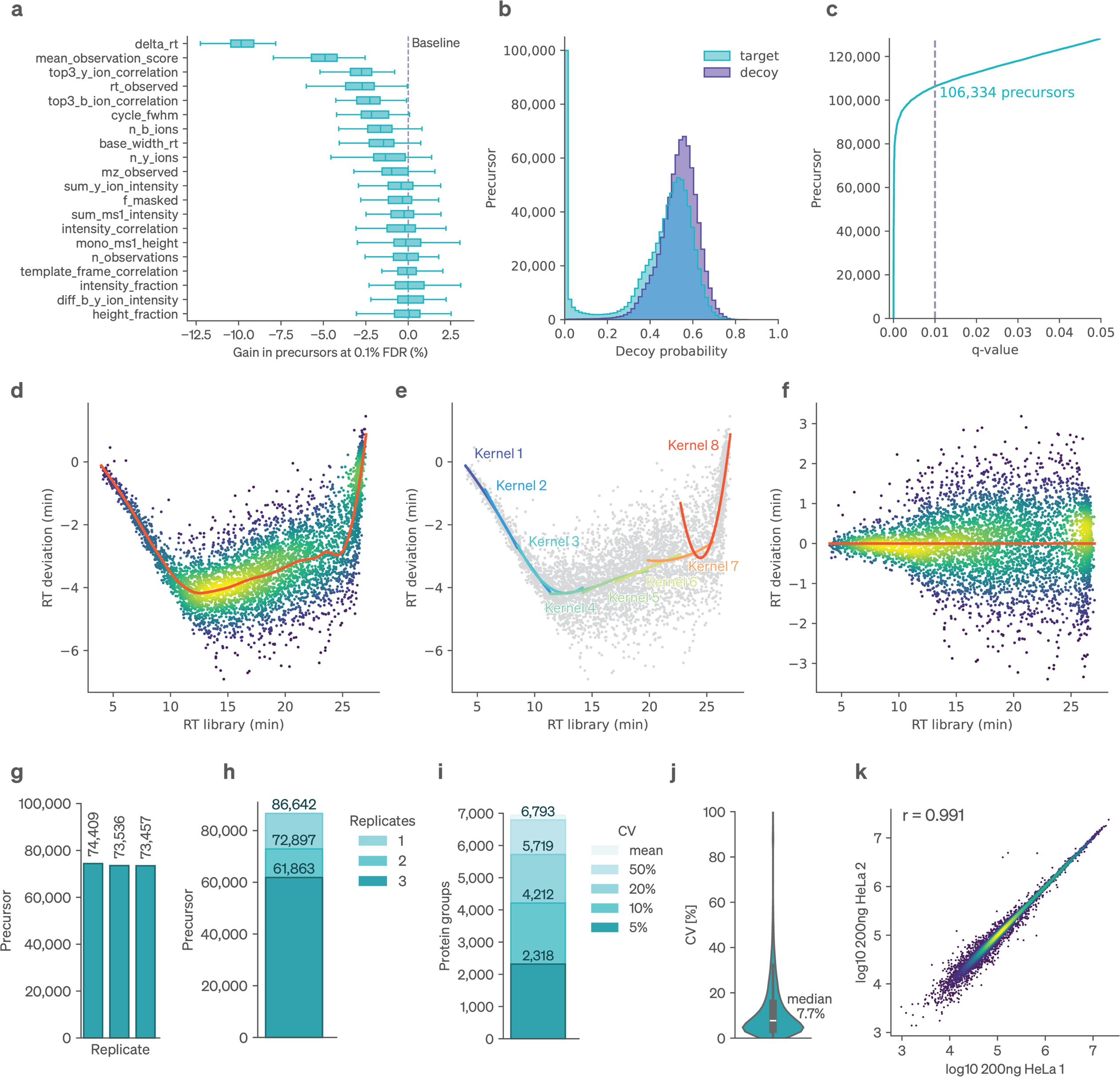
Central search engine components. a Classifier features and their importance for the supervised target decoy competition. **a**, Feature importance is defined as percentage drop of precursor identifications at 0.1% FDR. **b**, Deep neural network output probability for decoy peptides. **c**, Number of precursors identified as a function of the q-value cutoW. **d**, Non-linear calibration of retention times using LOESS regression (**Extended Data Fig. 3** and **Methods**). **e**, Collection of polynomial basis-functions combined using local kernels. **f**, Retention time deviation after calibration. **g-k**, Results for the library-based search of HeLa lysates measured with dia-PASEF. **g**, Number of precursors identified at a 1% FDR in three replicates. **h**, Precursors shared across replicates. **i**, Protein groups identified at given coeWicient of variants (CVs). **j**, Distribution of protein group CVs. **k**, Pearson correlation of precursor intensities across samples.

### AlphaDIA adapts to di=erent instruments and enables new acquisition methods

Recently, DIA has been coupled to sophisticated data acquisition schemes where the quadrupole isolation window scans nearly continuously through the m/z or m/z and ion mobility space^11,24,27^. The methods, termed synchro-PASEF or midia-PASEF hold the promise of much improved precursor specificity and quantitative accuracy, which, however, has been difficult to realize due to lack of flexible algorithms handling the thousands of individual isolation windows per DIA cycle. AlphaDIA’s processing algorithm and alphaRaw’s efficient data handling allows to use all synchro scans which contribute signal for a given precursor, considering its isotope distribution as a prior (**Fig. 3, a**). Using the masses and abundance of the precursor isotopes we model the behavior of the quadrupole, resulting in a template with the expected intensity distribution across synchro scan observations (**Fig. 3, b**). This template includes the slicing of the isotope distribution by the quadrupole which must be recapitulated in the intensity profiles of the fragments (**Fig. 3, c**). This comparison of the fragment profile with the template contributes to our deep-learning based identification score and enables analysis of complex proteomes (**Fig. 3, d, Extended Data Fig. 4**). This first processing algorithm for sliding quadrupole data could be extended from synchro-PASEF to similar acquisition schemes such as midia-PASEF or scanning SWATH.

**Fig. 3:**
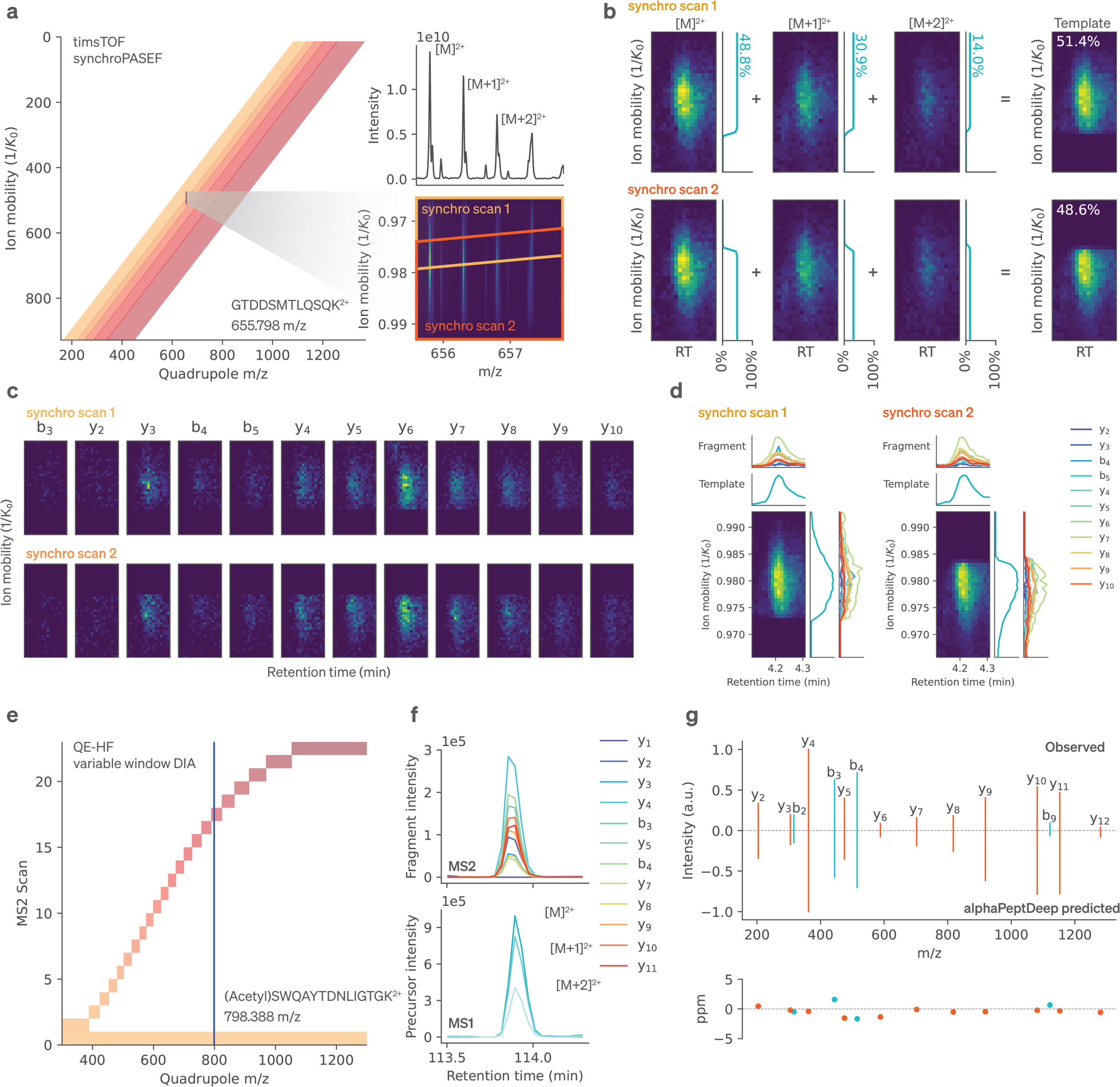
AlphaDIA enables flexible processing for diJerent acquisition methods. **a**, Variable window synchro-PASEF acquisition on the timsTOF. The quadrupole mass filter moves as precursors are released from the TIMS trap. The precursor with sequence GTDDSMTLQSQK is sliced by the quadrupole, resulting in fragment signal across two synchro scans. **b**, Slicing patterns are resolved by calculating the expected distribution of fragment signal in form of a template matrix. The template matrix is calculated by transforming the individual precursor isotope signal with the quadrupole transmission function of the synchro scans. **c**, Observed fragment signal across the two synchro scans. **d**, For each of the two synchro scans the elution and ion mobility XICs are compared. Comparison of the fragment signal (rainbow colors) to the template (blue) provides evidence of the identification of peptides. **e**, Application of the processing algorithm to variable window DIA data without ion mobility separation on a quadrupole Orbitrap analyzer (QE-HF). For the given precursor (Acetyl)SWQAYTDNLIGTGK all valid MS2 scans contributing evidence are selected. **f**, Elution profile of MS2 (top) and MS1 (bottom) ions for the precursor of interest. **g**, Observed and predicted fragment intensities after integration of the peak area (top) and mass accuracy for the same precursor (bottom).

Next, we wanted to extend the reach of alphaDIA to other proteomic platforms and methods. For instance, our algorithms adapted naturally to fixed as well as variable window DIA data from quadrupole Orbitrap analyzers. The absence of ion mobility reduces the search space to a one-dimensional search across retention time while still utilizing all valid MS2 observation for a given precursor (**Fig. 3, e**). As before, after discrete peak group candidates have been identified (**Fig. 3, f**) the spectrum centric view allows detailed scoring utilizing alphaPeptDeep predicted spectra (**Fig. 3, g**). Additionally, alphaDIA can process Orbitrap and Orbitrap Astral data with wide, narrow, variable or overlapping DIA windows. It can likewise process Sciex SWATH data (**Extended Data Fig. 5**).

### AlphaDIA matches or exceeds popular packages in empirical library-based search

Having established the ability of alphaDIA for in-depth analysis of complex proteomes and its adaptability to diverse platforms, we next wanted to directly benchmark its performance against other common DIA search engines. To avoid potential bias, we build upon a recently published benchmarking study from the Shui group, in which mouse brain membrane isolates were spiked into a complex background of yeast proteins in varying ratios and measured on a quadrupole orbitrap (QE-HF) and a timsTOF^33^. The authors generated empirical libraries with MS Fragger^4^ and optimized search parameters for DIA-NN, Spectronaut and MaxDIA (**Fig. 4, a**).

**Fig. 4:**
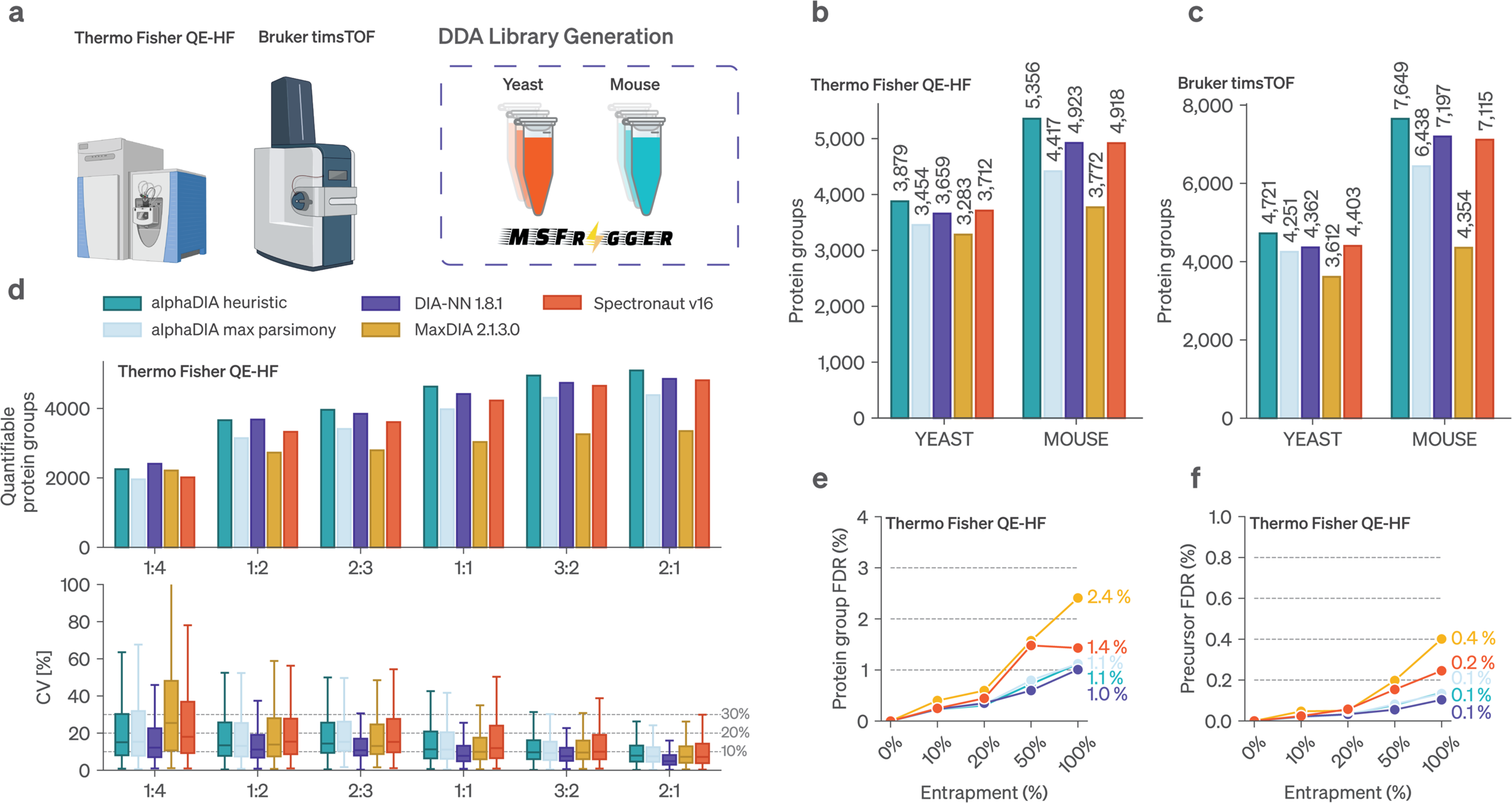
Benchmarking alphaDIA against established software for library bases DIA search. **a**, Overview of the benchmarking dataset^33^ for empirical library based search acquired on the quadrupole orbitrap QE-HF platform and the timsTOF. Fractionated bulk samples are analyzed using DDA to generate sample specific libraries using MSFragger. Mouse brain membrane isolates are spiked into a complex yeast background at diWerent ratios and analyzed in 5 replicates using DIA on both platforms. **b**, Number of Mouse protein groups identified at 1% FDR across all replicates on the QE-HF. **c**, Same as b but on the timsTOF platform. **d**, Quantified Mouse protein groups between diWerent spike ins and a reference sample. Proteins we’re deemed quantifiable if they were observed in at least 3 out of 5 replicates. The coeWicient of variation (CV) is shown for each set of identifications. **e**, Benchmarking of false discoveries using increasing amounts of Arabidopsis entrapments compared to the Yeast / Mouse spectral library. The false discovery rate on the protein level is shown for the QE-HF platform. **f**, Same as e, but on the precursor level.

Based on the provided libraries alphaDIA identified up to 50,600 mouse peptides in the QE data across all samples and up to 81,500 on the timsTOF (**Extended Data Fig. 6**). Inferring proteins from uniquely identified peptide involves considerations that can influence the number of reported protein groups^34^. AlphaDIA allows strict (maximum parsimony) or commonly used ‘heuristic’ grouping (**Methods**). With the latter, we identified 5,366 proteins (QE-HF) and 7,649 (timsTOF) protein groups across all samples, matching and even exceeding the other algorithms (**Fig. 4, b-c**). This is also reflected across replicates for single conditions. AlphaDIA quantified the most protein groups in at least 3 out 5 replicates for most ratios while maintaining comparable coefficients of variation (CV) and accuracy as judged by the proteome mixing ratios (**Fig. 4, d, Extended Data Fig. 6-9**).

To prevent over-reporting by sophisticated DIA database searching strategies based on internal target decoy FDR estimates, results can be externally validated by including additional proteome databases from species not present in the sample^35^. As in the benchmarking study, we performed an entrapment search with an Arabidopsis library added in increasing proportions to the target library. On both MS platforms, even for 100% entrapment Arabidopsis identifications matched the chosen target FDR of 1% at the protein level **Fig. 4**, e-f). At this protein FDR, false positive precursors are even less likely appearing only at 0.1% globally. This contrasted with some of the other tested tools, which reported up to three-fold more false positive Arabidopsis identifications than intended at the chosen FDR target (**Extended Data Fig. 8, a-d**). Importantly, the increased library size only minimally decreased overall identifications for alphaDIA (**Extended Data Fig. 8, e-h**). We conclude that for library-based search alphaDIA provides at least competitive performance with common search engines while maintaining a reliable and conservative FDR.

### Combining alphaDIA and alphaPeptDeep allows fast search of fully predicted libraries

While empirical libraries benefit from implicitly capturing instrument and workflow specific properties, the key advantage of deep-learning predicted libraries of the entire proteome database is that it eliminates cumbersome library measurement altogether. We recently introduced alphaPeptDeep, an open source, transformer-based deep learning framework for predicting all MS-relevant peptide properties from their sequences^20^.

With these state-of-the art predicted libraries, we devised a two-step search workflow in alphaDIA consisting of library refinement and quantification (**Fig. 5 a**). Furthermore, we reasoned that our feature-free search should adapt well to the high sensitivity TOF data generated by the Orbitrap Astral mass spectrometer. For benchmarking, we acquired and searched bulk Hela samples with an alphaPeptDeep predicted library containing 3.6 million tryptic precursors. AlphaDIA identified on average more than 120.000 precursors, matching or exceeding the performance of all other tested search engines (**Fig. 5 b**). Remarkably, in this 60 SPD method (21 min) this corresponded to the identification of 9,500 protein groups of which 8,200 had a CV less than 20% (**Fig. 5 d**). The great depth of proteome characterization was also reflected in the data completeness across replicates (**Extended Data Fig. 10**). Search times stayed below the rapid acquisition time (**Fig. 5 e**). We validated the FDR control of this more complex two step workflow using the entire Arabidopsis library, which externally confirmed rigorous control of false positive identifications (1.08% at protein level and 0.2% at precursor level, **Fig. 5, f**).

**Fig. 5:**
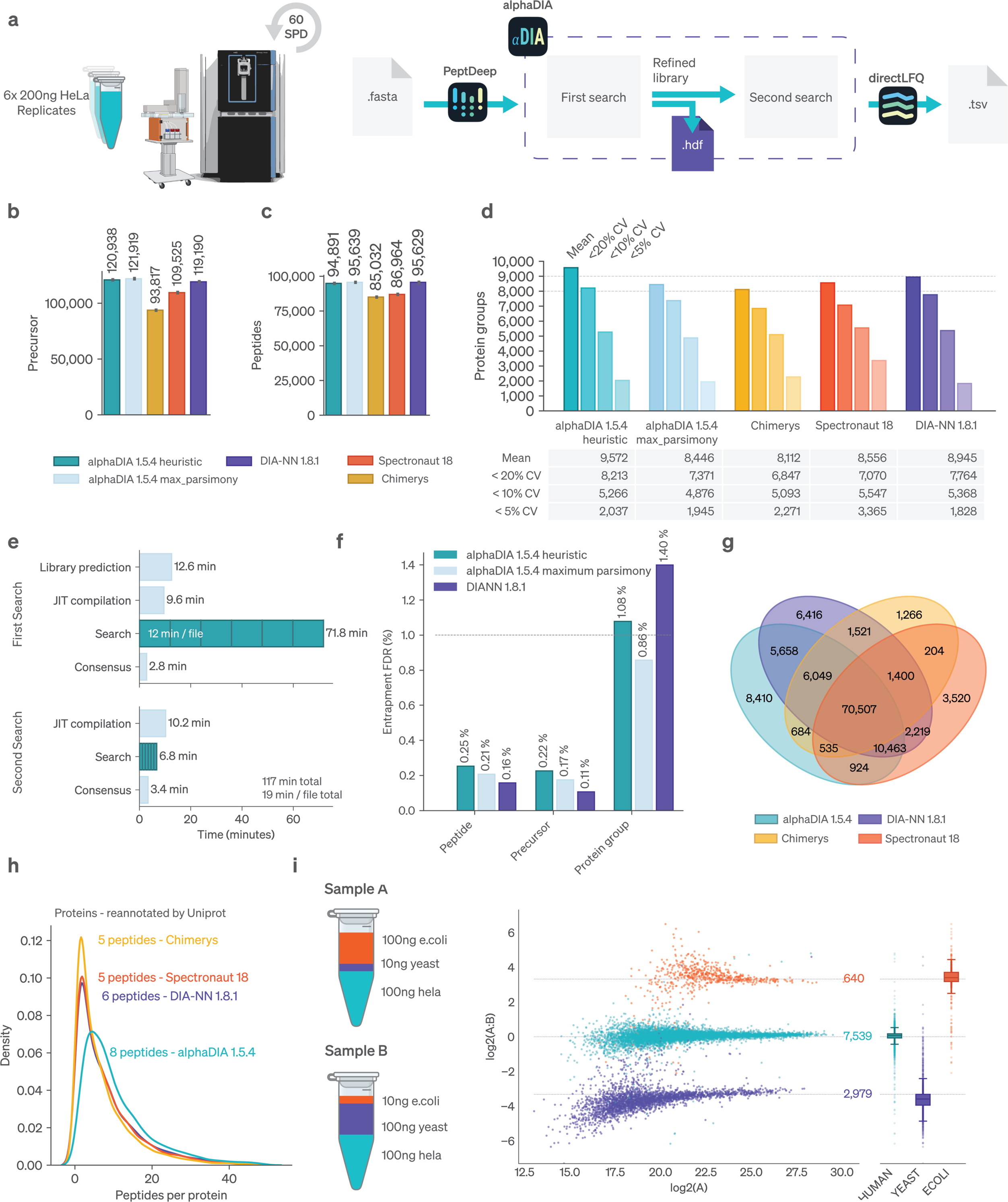
Searching complex proteomes acquired on the Orbitrap Astral with fully predicted spectral libraries. **a**, Six replicates of 200ng HeLa bulk data were analyzed on the Orbitrap Astral with a 60SPD (21 min) gradient. A fully predicted alphaPeptDeep library was used for a two-step search in alphaDIA. DiWerent search engines were used for comparison. **b**, Mean precursors identified across search engines **c**, Mean protein groups identified across processing methods **d**, Protein groups identified at given coeWicients of variation (CV) cutoWs. **e**, Analysis time for diWerent processing steps when analyzed with on a 32 core machine. **f**, Arabidopsis entrapment search using the fully predicted library workflow. The share of identified Arabidopsis proteins at 1% target decoy FDR is shown. **g**, Venn diagram showing the overlap of proteotypic peptides across processing methods. **h**, Analysis of protein overlap between diWerent processing methods. Peptides were mapped back to the same reference proteome, discarding ambiguous matches. Number of peptides identified per protein. The median number of peptides per protein is shown. **i**, Mixed species experiment for establishing quantitative accuracy. Human, Yeast and E.coli proteomes were combined in defined ratios. Plotting the ratio between species-unique protein groups recapitulates the expected ratio (dashed lines).

To compare identified proteins across search engines, we mapped peptide sequences to the UNIPROT reference proteome, discarding ambiguous peptides mapping to multiple proteins. Reassuringly, more than 70,000 peptides and close to 8,000 proteins were jointly identified by all tested tools (**Fig. 5 g**). AlphaDIA had the highest number of uniquely identified peptides among search engines, manifesting in higher sequence coverage (median of 8 peptides per protein, **Fig. 5 h**).

To assess the accuracy of label-free quantification (LFQ), we used the established strategy^36^ of three species proteomes mixed in defined ratios, acquired on the Orbitrap Astral. Fully predicted library search combined with directLFQ recapitulated the expected ratios with excellent precision and accuracy (**Fig. 5 i, Extended Data Fig. 11**).

Multiplexed DIA has recently shown great potential to increase throughput and depth^37,38^. To analyze such data, identifications must be transferred between the channels which involves an additional channel FDR. Due to the modular nature of alphaDIA this functionality was readily incorporated. We benchmarked it on a DIA dataset in which HeLa cells were heavy and light SILAC labeled and analyzed on a QE-HFX^39^ (**Extended Data Fig. 12**). In proportions of identifications in ‘light only’, ‘heavy only’ and ‘light and heavy’ were very similar to the previous DDA and DIA results, validating our channel FDR. Interestingly, on the same data the absolute number of identified peptides was threefold higher than in the original paper, reflecting advances in DIA search over the last years in general, and specifically in alphaDIA.

### DIA transfer learning generalizes DIA search to unseen modifications

To date, fully predicted libraries address many of the needs of DIA workflows but their pretrained prediction models are still best suited to the sample and instrument types that were used in training. This makes it necessary to train custom models for different situations - for example PTMs, as they generally change retention and fragmentation behavior compared to the unmodified peptide. We reasoned that close integration of prediction by deep learning and the search engine might have the potential learn to adapt to such differences, an approach that we call *end-to-end transfer learning*. Following search with alphaDIA confidently identified precursors and their spectra are first collected into a training data set. The general pretrained models for retention time, fragmentation spectra and charge state provided with alphaPeptDeep are then finetuned using transfer learning on the experiment specific training data set (**Fig. 6, a, b**). This results in a custom model, reflecting the behavior of peptides on the individual LCMS setup. A held-out test data set ensures generalization and prevents overfitting.

**Fig. 6:**
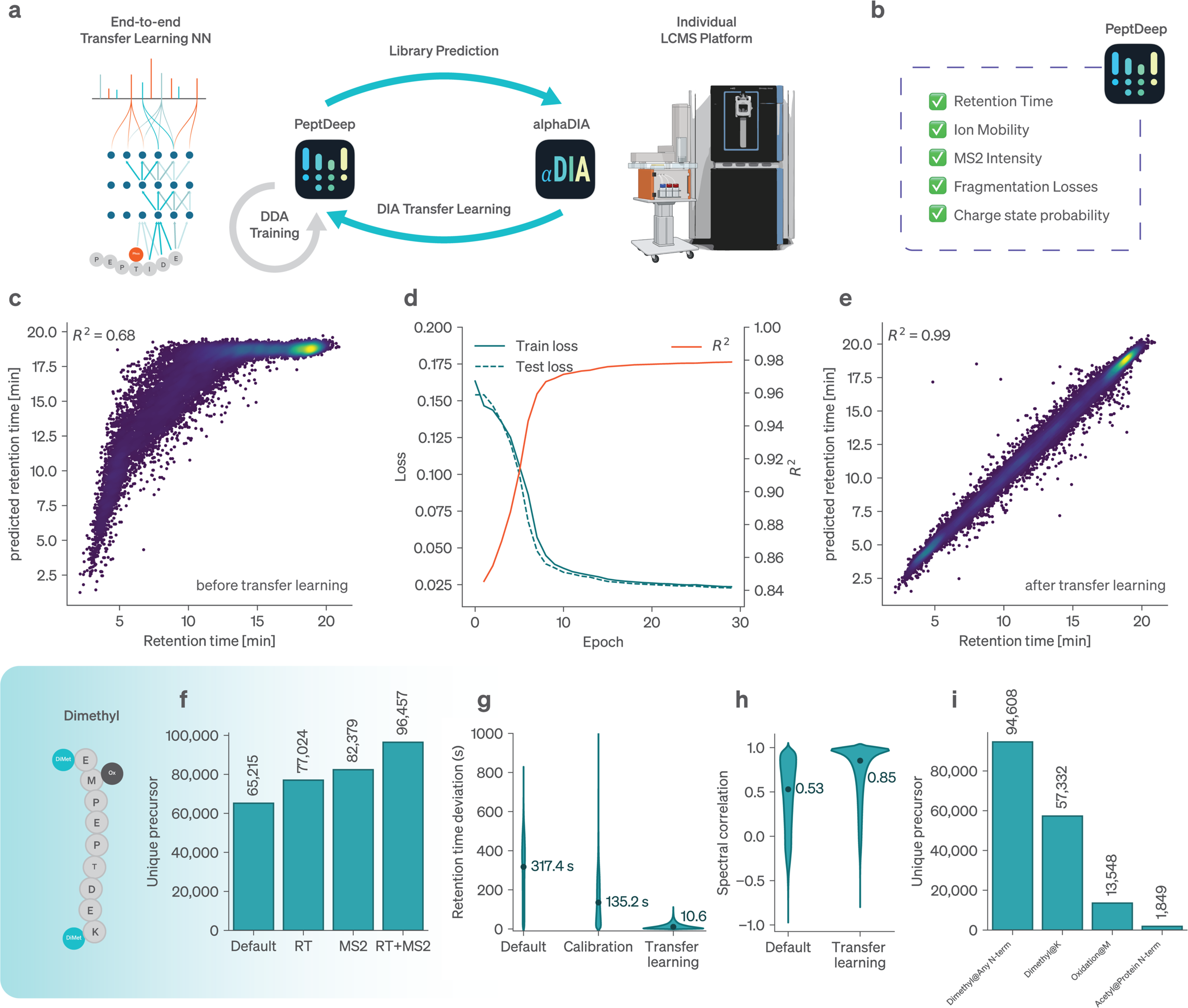
DIA transfer learning for discovery of modified peptides. **a**, A custom deep learning model is trained for every experiment using the identifications from the DIA search engine. b, Multiple properties are being optimized resulting in smaller and better matching spectral libraries. c, Observed and predicted retention times for dimethylated precursors before transfer learning. d, DIA transfer learning for the retention times of dimethylated peptides. During training by stochastic gradient descent, a 20% test set of precursors is held out to mitigate overfitting and ensure generalization to the peptide space of interest. e, Retention times after transfer learning. f, Comparison of the number of unique peptides identified with the pretrained base model (Default) to the transfer learned model after RT and MS2 transfer learning. g, Distribution of absolute retention time errors for the pre trained base model (Default), the non-linear calibration within alphaDIA and after transfer learning. h, Comparison of spectral correlation before and after MS2 transfer learning. i, Number of unique observed modifications by type.

To assess the potential of this end-to-end transfer learning concept, we first applied it to a dataset of dimethylated HeLa peptides, an example of a modification that is known to alter retention times and fragmentation behavior (**Methods, Fig. 6, c**). We found that transfer learning accurately modeled the effects of the lysine and N-terminal dimethylation on retention time behavior, improving R^2^ from 0.69 to 0.99 (**Fig. 6**, d-i).

Using the transfer learned model resulted in a total of 96,000 unique precursor and 8,613 protein identifications, a 48% increase over the 65,000 precursors identified without transfer learning and a 25% increase in protein groups (**Fig. 6, d,e**; **Extended Data Fig. 14**). This gain in identifications is driven additively by both improved predictions of retention times from a median prediction error of 317 s down to only 11 s and an increase in the median correlation to predicted spectra from 0.5 to 0.85 (**Fig. 6, g,h**).

Given these drastic improvements, we wished to ascertain that they were not the result of overfitting, despite the use of a holdout test dataset. Similarly to before, we used entrapment with the Arabidopsis proteome library followed by transfer learning with all precursors, including false positive Arabidopsis hits (**Extended Data Fig. 13,a**). Remarkably, even successive rounds of transfer learning led to more confident precursors identifications and less than 0.5% false Arabidopsis identifications at 1% FDR (**Extended Data Fig. 13, b-d**). Upon inspection, we found that predictions of target hits showed substantial improved agreement with observed data, whereas the opposite was true of false positive Arabidopsis hits (**Extended Data Fig. 13, e-g**). This implies that end to end transfer learning generalizes to the peptide behavior in the actual experiment improving identifications and control of false discoveries at the same time.

## Discussion

The development of alphaDIA addresses several critical challenges inherent to DIA, such as the complexity of spectral data and the need for robust, adaptable algorithms capable of handling high-dimensional data from advanced instrumentation. Our results demonstrate that already the first public version of alphaDIA matches and in many cases surpasses existing software tools in terms of performance and versatility, making it a valuable addition to the proteomics toolkit.

AlphaDIA’s feature-free processing method is central to its performance and flexibility. Traditional DIA processing methods often rely on predefined feature boundaries, which can lead to information loss, especially with the high sensitivity and stochastic nature of TOF detectors. By contrast, alphaDIA’s approach aggregates signals across multiple dimensions, ensuring that all relevant data is utilized before making discrete identifications. This results in higher accuracy and sensitivity, as evidenced by our ability to confidently identify peptides even in noisy datasets. Additionally, alphaDIA extends the reach of DIA to novel acquisition modes. Together with its open-source architecture this enables the community to quickly loop between experimental innovations and their algorithmic implementation.

Our benchmarking against established tools using both empirical and predicted libraries showcases alphaDIA’s equal or superior performance. This holds true across platforms and experimental designs including the Orbitrap Astral, where alphaDIA identified over 120,000 precursors and 9,500 protein groups in a 60 SPD format.

One of the most innovative and promising aspects of alphaDIA is its end-to-end transfer learning capability. Based on integration with the transformer models of alphaPeptDeep, alphaDIA closes the loop between spectral library prediction and DIA search. Our approach allows the model to adapt to experiment-specific conditions, enhancing the accuracy of peptide identifications. We showcased this on a dataset of dimethylated HeLa peptides demonstrating dramatic improvements in retention time prediction and spectral correlation, resulting in a 48% increase in unique precursor identifications and a 25% increase in protein groups compared to using pretrained models alone. This allows the application of DIA search to hitherto inaccessible areas such as post-translationally modified proteins without PTM specific pretraining or to the better identification of HLA peptides. Importantly we demonstrated that transfer learning not only improves overall identifications but even improves FDR control, ensuring reliable results.

The advancements presented by alphaDIA pave the way for more comprehensive and accurate proteomic analyses which will be important as MS technology continues to evolve. This will be especially important in clinical and translational research, where ever increasing cohorts and data require large scale, distributed processing.

The framework’s open-source nature ensures that it can be continuously improved and extended by the scientific community, fostering innovation and collaboration. We therefore aim to establish alphaDIA as a cornerstone for the next generation of DIA analysis, closely coupled to the developments in artificial intelligence.

## Acknowledgments

We thank Mann Labs members and Isabell Bludau for insightful discussions. This work is funded by the Max Planck Society for the Advancement of Science, and by the Bavarian State Ministry of Health and Care through the research project DigiMed Bayern (www.digimed-bayern.de). It was supported by European Union’s Horizon 2020 research and innovation program under grant agreement No. 874839 (ISLET).

## Contributions

Conceptualization: G.W., WF.Z and M.M. Bioinformatic method development G.W., M.L., V.B., M.K., C.A., WF.Z. Architecture of ecosystem algorithms & software WF.Z., C.A., G.W., M.K., M. Sch., M.St. S.W. Proteomics method development and data acquisition T.H., P.S., M.T., S.S., Writing - original draft: G.W. and M.M. Writing - review and editing: all authors; Resources: all authors. Supervision: M.M.; Funding acquisition: M.M.

## Code Availability

All code presented herein as part of alphaDIA is free software accessible under the permissive Apache license. **AlphaDIA** can be found at www.github.com/MannLabs/alphadia, **alphaRaw** can be found at www.github.com/MannLabs/alpharaw, **alphaBase** is found at www.github.com/MannLabs/alphabase.

## Data Availability

All data will be made available upon publication of the manuscript.

## Methods

### Calibration of retention time, ion mobility and m/z

During search retention time, ion mobility, precursor m/z and fragment m/z are calibrated to the measured values. Starting with initial default settings of 15ppm MS1 and MS2 tolerance, 300 seconds rt tolerance and 0.04 mobility tolerance the library is iteratively calibrated within a minimum of three epochs. Every epoch, batches of precursors are searched and scored with an exponential batch plan (2000, 4000, 8000, etc.) until a minimum number of precursors has been identified at 1% FDR. The number of target precursors is increase with every epoch (default: 200 precursors/epoch). If one epoch has accumulated enough confident target precursors, they are calibrated to the measured values using locally estimated scatterplot smoothing (LOESS) regression. For calibration of fragment m/z values, up to 5000 (but at least 500) of the best fragments according to their XIC correlation are used. Following a single calibration pass, all tolerances are updated to the 95 percentile error after calibration but not below the chosen target level.

LOESS regression using uniformly distributed kernels is used for each property which should be calibrated (**Extended Data Fig. 3**). Regression is performed on first and second degree polynomials basis functions of the calibratable property. For m/z and ion mobility, two local estimators with tricubic kernels are used. For retention time prediction, six estimators with tricubic kernels are used. The architecture is built on the scikit-learn package and can be configured to use different hyperparameters and arbitrary predictors for calibration.

### Scoring of precursors and decoys using convolution kernels and supervised classification

AlphaDIA employs a two-step scoring machine learning algorithm to identify the best potential peak group for every library entry. The first step builds on a collection of weighted convolution kernels, learned during optimization and calibration of the spectral library. For every precursor of interest, MS1 scans and MS2 scans contributing information towards the identification are identified from the DIA cycle pattern of the acquisition method. Based on a certain number of highest intensity fragments in the library (default: 12), dense representations of the search space in ion mobility and retention time dimension are assembled. To identify putative peak groups for each precursor, a set of convolution kernels, reflecting the expected distribution in retention time, ion mobility and fragment intensity are learned during calibration and optimization. The convolution of the search space is performed in Fourier space for fast processing, and a single score is calculated as log sum across kernels and fragments. Local maxima are identified using a simple peak picking algorithm and retention time and ion mobility boundaries of the peak group of interest are defined from the joint scoring function. These candidates are subsequently rescored for FDR estimation.

As second step, AlphaDIA uses target decoy competition for scoring the quality of precursor spectrum matches. Upon library import, paired known false positive decoy peptides are created for every target. By default, a mutation pattern GAVLIFMPWSCTYHKRQENDBJOUXZ => LLLVVLLLLTSSSSLLNDQEVVVVVV is used. For every library entry, target and decoy, the best high scoring matches from the convolution kernel score are used for supervised classification. Up to 47 features are calculated for each peak-group match, reflecting the merit of the identification. A multi-layer perceptron (MLP) deep neural network with layer sizes 100, 50, 20, 5 and 47 input dimensions (10,810 parameters) is trained to predict the probability of being a false decoy identification. Training is performed with stochastic gradient descent for 10 epochs with a batch size of 5000 and learning rate of 0.001. While training on an 80% training set a 20% test set is held-out to mitigate overfitting. Based on the final score, the best (lowest) decoy probability peak group is retained for every library entry and a count based FDR is calculated.

### False discovery rate calculation

AlphaDIA uses a count based FDR on the level for assigning confidence to precursor, peptide, protein and channels. Identifications are given as a set of target and decoy identifications *P* = {*p*_0_, *p*_1_,…*p*_i_} all associated with a ground truth decoy status *decoy*: *P* → {*true*, *false*} and a deep-learning derived decoy score 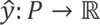. For every precursor with index *i* the number of targets with lower or equal decoy probability

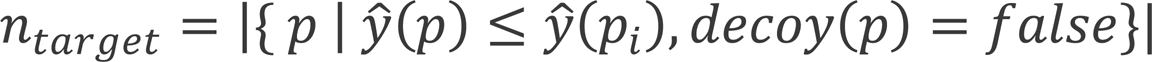

and the number of decoys with lower or equal decoy probability

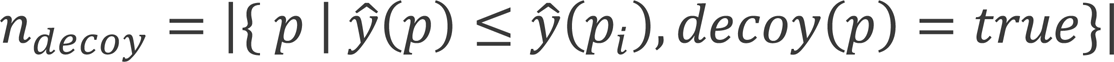

are calculated. Furthermore, the total number of targets and decoys in the set are calculated as:

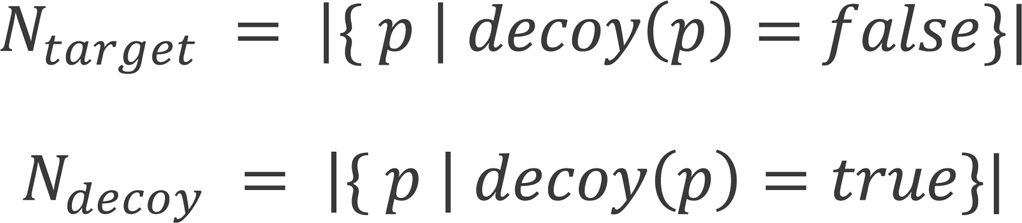

The local count-based q value is given as:

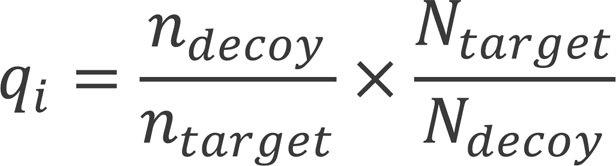

This is converted to a false discovery rate (FDR) by using the minimum q-value where a precursor was accepted:

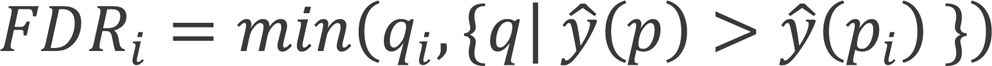

By default, all identifications are filtered on a run-level 1% FDR precursor threshold and global 1% protein group-level threshold.

### Spectrum centric fragment competition

Competition of precursors for fragment ion is used as spectrum centric element to mitigate double use of fragments for multiple identifications from the same spectra. Following initial FDR calculation, precursor candidates are filtered at 5% FDR and split into groups of potentially fragment sharing. This is determined by the quadrupole cycle pattern. Then, precursor candidates and their elution width at half maximum are compared so that precursors with overlapping elution width at half maximum have no more than *k_max_* = 1 shared fragment masses within the chosen MS2 mass accuracy δ_*MS*2_. If two or more precursor candidates share more fragments than permitted the precursor candidate with the lowest decoy score is used.

### Protein inference

Reporting all proteins whose sequence can be matched to any identified peptide can lead to drastic inflation of false discoveries on the protein level^40^. Following the approach outlined by Nesvizhskii et al. ^41^, we consider a precursor as a single piece of evidence, and the task of protein inference is then to assemble these precursors into proteins while controlling the accumulation of spurious protein identifications. AlphaDIA aims to implement a simple and transparent inference approach, allowing for three inference modes: library, maximum_parsimony and heuristic. Apart from the library mode which uses the inference performed during empirical library creation, protein inference is based on an implementation of the “greedy set cover” algorithm with grouping by default (heuristic) and without grouping for strict inference (maximum_parsimony).

In brief, alphaDIA’s protein inference starts with a table of identified precursors. Each precursor is associated with a set of genes and proteins and based on user choice, the inference is performed on the gene or protein level (default: gene). While a common peptide precursor may match many proteins, a proteotypic peptide will match one single protein. During grouping, the precursor and protein arrays are reshaped into a protein-centric view, where each protein is associated with one set of precursors. Then, proteins are sorted by the length of their precursor set in descending order, and the protein with the largest number of precursors removed from the lists as the first query. The query is compared to all remaining subject proteins. From each subject precursor set, all precursors matching the query set are removed. If a protein’s precursor set becomes empty, it is considered redundant and dropped. After all precursor sets have been compared, the process repeats by reordering the list and extracting the next query. After completion, retained queries are denoted master proteins, necessary to explain all discovered precursors. In strict maximum_parsimony mode all master proteins are simply reshaped to precursor-centric format, linking each precursor to one single protein ID. In the heuristic mode, the list of master proteins is used to remove all non-master proteins from the initial precursor table, effectively leaving each precursor with a set of associated proteins comprised solely of master proteins. Thereby, the same precursor can be claimed by different proteins, creating protein groups (see also the tutorial notebook in the GitHub repository).

### Protein FDR

Protein FDR is performed on the protein groups (PGs) calculated during protein inference. For all target and decoy protein groups, 7 features are calculated: the total number of precursors across runs for the PG; the mean decoy score for precursors across runs for the PG; the number of unique peptides for the PG; the number of unique precursors for the PG; the number of runs the PG was found in; the lowest decoy score across precursors for the PG; the highest decoy score across precursors for the PG. We use a multi-layer-perceptron (MLP) to classify decoy PGs from target PGs. Correct training is ensured by a 20% held-out test set. PG FDRs are calculated on a global level using the FDR mechanism described just above.

### Library refinement for fully predicted libraries

AlphaDIA uses an established two step-search strategy for library refinement^15^. Following an initial search of all or a subset of raw files, protein inference and FDR is performed as configured by the user. All precursors are automatically filtered at 1% local precursor FDR and global 1% protein group FDR and accumulated into a spectral library and finally saved to the project folder. For each precursor, the identification with the best (lowest) decoy probability is used. By default, MS2 quantities are used as annotated in the original library. If transfer learning accumulation is used, custom user specified fragment types can be selected and observed MS2 intensities are extracted. This spectral library is then used for the second search with full MS2-based target decoy scoring without any relaxed FDR parameters. For protein inference and FDR, library annotated protein groups are used.

### Transfer learning

To create transfer learning libraries, precursors identified at 1% precursor and protein FDR are selected for requantification. Precursors are requantified for user defined fragment ion types (a, b, c, x, y, z, modification loss, etc.) and a user-defined maximum charge (default: 2). Extracted fragment quantities are accumulated across samples and ordered by their decoy probability. For each unique modified precursor, the observations with the three lowest decoy scores are selected. AlphaDIA also creates a high quality subset where only precursors with a median fragment correlation greater than 0.5 are included. For these precursors we only retain fragments whose correlation values exceed 75% of the median fragment correlation of the respective precursor. The implementation of transfer learning library is globally sequential. At any given time, we can limit the implementation to only parallelize across a limited number of processes. This approach allows the process to scale without storing all runs in memory.

For transfer learning, we prioritized robustness to ensure performance instead of requiring users to define hyperparameters. The transfer learning dataset is split into a training (80%) and test set (20%) and trained for a maximum of 50 epochs. After each training epoch, we run a test epoch for assessing the test loss and data specific test metrics. AlphaDIA uses a custom learning rate scheduler with two phases. The first phase is a warm-up period (default 5 epochs) during which the learning rate gradually increases to a maximum value (default: 0.005). After this warm-up phase the learning rate scheduler halves the learning rate if the training loss does not significantly improve (default: >5% test loss) within a patience period (default: 3 epochs). Additionally, we use a simple early stopping mechanism that interrupts training if the validation loss starts to diverge or does not significantly improve (default: 12 epochs).

After training, the deep learning model is stored on disk, and can be loaded as necessary. Retention time and ion mobility finetuning are supervised by calculating the L1 loss, R2, 95th percentile of the absolute error on the training data. MS2 finetuning is supervised by calculating the L1 loss, Pearson correlation coefficient, spectral angle, Spearman correlation on the test data. Charge finetuning is supervised by calculating the cross entropy loss, accuracy, precision, recall on the test data. All training and test metrics are reported to the user. The specific implementation and details of the test metrics can be found in the open-source code on GitHub (see **Code Availability**).

### Sample preparation of HeLa bulk digests

HeLa S3 cells (ATCC) were cultured in Dulbecco’s modified Eagle’s medium (Life Technologies Ltd) supplemented with 20 mM glutamine, 10% fetal bovine serum, and 1% penicillin-streptomycin. After washing the cells in PBS and cell lysis, the proteins were reduced, alkylated, and digested by trypsin (Sigma-Aldrich) and LysC (WAKO) (1:100, enzyme/protein, w/w) in one step. The peptides were dried, resuspended in 0.1% TFA/2% acetonitrile (ACN), and 200 ng digest was loaded onto Evotips (Evosep). The Evotips were prepared by activation with 1-propanol, washed with 0.1% formic acid (FA)/99.9% ACN, and equilibrated with 0.1% FA. After loading the samples, tips were washed once with 0.1% FA.

### Sample preparation of dimethylated peptides for transfer learning

HeLa cells were cultured as describe above. A HeLa cell pellet was lysed by boiling for 10 min in 1 % SDC in 60 mM TEAB pH 8.5, followed by sonication in a Branson type instrument, Heinemann Sonifier 250 (Schwäbisch Gmünd), operating at 20% duty cycle and 3-4 output for 1 min, and boiling for 5 min again. After cooling down to room temperature, the protein concentration was determined using the tryptophan fluorescence based, WF-assay in the microtiter plate format using white Nunc 96-well plates with a flat bottom (Thermo Fisher Scientific, 136101). After diluting the lysate to 1 ug/uL in lysis buffer, disulfide bonds were reduced by adding Tris(2-carboxyethyl)phosphine (TCEP) to a final concentration of 10 mM TCEP and briefly incubating for 10 min. Denatured protein lysate was digested by Arg-C Ultra (Promega) and Lys-C (WAKO) at a 1:250 and 1:100 (enzyme/protein) ratio to the lysate at 37°C for 3 h, respectively. The peptides were labeled with a dimethyl group by using a 100 uL of 1 ug/uL digested peptides and adding 4 uL of 4 % formaldehyde and 4 uL of a 0.6 M NaBH3CN solution. The mixture was incubated at room temperature and every 10 minutes 2.8 uL (2 ug peptides) were sampled until 60 minutes and added to 17.2 uL of a 1 % solution of trifluoro acetic acid to quench the reaction.

### Sample preparation for the mixed species experiments

For the mixed species experiment, three different mixtures with varying mixing ratios of HeLa tryptic digest (Pierce #1862824), S. cerevisiae tryptic digest (Promega V746A), and E. coli tryptic digest (Waters #186003196) were prepared: Sample A (10:1:10 Human(H):Yeast(Y):E. coli(E)), Sample B (10:10:1 H:Y:E), and Sample C (10:4:7 H:Y:E). Five replicates containing 210 ng were loaded per condition.

### Peptide loading onto C-18 tips

C-18 tips (Evotip Pure, Evosep) were loaded with the Bravo robot (Agilent), by activation with 1-propanol, washing two times with 50 μl buffer B (99.9% ACN, 0.1% FA), activation with 1-propanol and two wash steps with 50 μl buffer A (99.9% H2O, 0.1% FA). In between, Evotips were spun at 700 g for 1 min. For sample loading, Evotips were prepared with 70 μl buffer A and a short spin at 700 g. Samples were loaded in 20 μl with the indicated concentration into the remaining buffer A and spun at 700 g for 1 min, if not described differently. After sample loading, Evotips were washed with 50 μl buffer A and stored with 150 μl buffer A after a short spin at 700 g at 4 °C until MS acquisition.

### MS data acquisition of dia-PASEF and synchro-PASEF data

We used the Evosep One liquid chromatography system to separate peptide mixtures at varying throughputs using standardized gradients. These gradients consisted of 0.1% formic acid (FA) and 99.9% water (v/v), and 0.1% FA with 99.9% acetonitrile (v/v) as mobile phases. For the 60 SPD runs, peptides were separated on a Pepsep column (8 cm x 150 μm ID, 1.5 μm C18, Bruker Daltoniks) connected to a 10 μm ID fused silica emitter (Bruker Daltoniks). For the whisper40 SPD runs, we utilized an Aurora Elite nanoflow column (15 cm x 75 μm ID, 1.7 μm C18, IonOpticks).

The system was coupled with a timsTOF mass spectrometer (Bruker Daltoniks) to acquire data in dia-PASEF and synchro-PASEF modes. Sample loads above 25 ng were analyzed using a timsTOF Pro2, and those below 25 ng with a timsTOF Ultra. The dia-PASEF and synchro-PASEF methods were optimized using our Python tool, py_diAID^32^. This tool maximizes precursor coverage by optimally positioning the acquisition scheme over the precursor cloud and enhances sampling efficiency by adjusting the isolation window widths according to precursor density.

The dia-PASEF method covers an m/z range from 300 to 1200 with eight dia-PASEF scans and two isolation window positions per scan (cycle time 0.98 s). The synchro-PASEF method covers an m/z range from 140 to 1350 with four diagonal synchro scans (cycle time 0.53 s). The method files are deposited in the data repository. In both modes, the fragment scans were acquired with an m/z range from 100 to 1700. Furthermore, ions were accumulated and ejected at 100 ms intervals from the TIMS tunnel. The methods cover an ion mobility range from 1.3 to 0.7 V cm^−2, calibrated with Agilent ESI Tuning Mix ions (m/z, 1/K₀: 622.02, 0.98 V cm^−2; 922.01, 1.19 V cm^−2; 1221.99, 1.38 V cm^−2). The collision energy was linearly decreased in relation to the ion mobility elution: from 59 eV at an ion mobility of 1.6 Vs cm^−2 to 20 eV at 0.6 V cm^−2.

### MS data acquisition of SWATH data on the ScieX 7600

Triplicates of 200ng HeLa bulk digest were loaded onto C-18 tips as described above and analysed using an Evosep One system (Evosep) coupled to a 7600 ZenoTOF mass spectrometer (Sciex) using Sciex OS (version 3.3 or higher). Peptides were separated by the 60 SPD method gradient (Evosep) on a PepSep 8cm x 150 μm reverse-phase column packed with 1.5 μm C18-beads (Bruker Daltonics) at 50 °C connected to the low micro electrode for 1-10 μL/min. The mobile phases were 0.1% formic acid in LC–MS-grade water (buffer A) and 99.9% ACN/0.1% FA (buffer B). The ZenoTOF mass spectrometer was equipped with the Optiflow ion source using a spray voltage of 4.5 kV, ion source gas 1 of 15 psi, ion source gas 2 of 60 psi, curtain gas of 35 psi, CAD gas of 7 and a temperature of 200 °C. SWATH data was acquired using the following parameters: TOF MS start mass of 400 Da, a stop mass of 1500 Da, TOF MS accumulation time of 50 ms, TOF MSMS start mass 140 Da, stop mass 1750 Da, accumulation time 13 ms with dynamic collision energy turned on, a charge state of 2, Zeno pulsing enabled, and 60 variable SWATH windows covering the mass range of 400-900 m/z.

### MS data acquisition of mixed species samples fostering innova-on and collabora-on on the Orbitrap Astral

For mixed species experiments, five replicates of samples A, B and C were loaded onto C-18 tips as described above. Samples were analyzed using an Evosep One system (Evosep) coupled to a Orbitrap Astral mass spectrometer (Thermo Scientific) using Thermo Tune software (version 1.0 or higher). Peptides were separated by the 60SPD method gradient (Evosep) on a PepSep 8 cm × 150 μm reverse-phase column packed with 1.5 μm C18-beads (Bruker Daltonics) at 50 °C. The analytical column was connected to a stainless-steel emitter with inner diameter of 30 µm (EV1086). The mobile phases were 0.1% formic acid in LC–MS-grade water (buffer A) and 99.9% ACN/0.1% FA (buffer B). The Orbitrap Astral mass spectrometer was equipped with a FAIMS Pro interface and an EASY-Spray source (both Thermo Scientific). A compensation voltage of −40V and a total carrier gas flow of 3.5 L/min was used as well as an electrospray voltage of 2.0 kV was applied for ionization. The MS1 spectra was recorded using the Orbitrap analyzer at 120k resolution from m/z 380-980 using an automatic gain control (AGC) target of 500% and a maximum injection time of 3 ms. The Astral analyzer was used for MS/MS scans in data-independent mode with 3 Th non-overlapping isolation windows with a scan range of 150-2000 m/z. The precursor accumulation time was 3ms and an AGC target of 500%. The isolated ions were fragmented using HCD with 25% normalized collision energy.

### MS data acquisition of HeLa bulk data on the Orbitrap Astral

For analysis of HeLa bulk digest, 200ng of lysate was loaded onto C-18 tips in six replicates as described above. Samples were analyzed using an Evosep One system (Evosep) coupled to a Orbitrap Astral mass spectrometer (Thermo Scientific) using Thermo Tune software (version 1.0 or higher). Peptides were separated by the 60SPD method gradient (Evosep) on an Aurora Rapid 80 mm × 0.15 mm reverse-phase column packed with 1.7 μm C18-beads (IonOpticks) at 50 °C. The mobile phases were 0.1% formic acid in LC–MS-grade water (buffer A) and 99.9% ACN/0.1% FA (buffer B). The Orbitrap Astral mass spectrometer was equipped with a FAIMS Pro interface and an EASY-Spray source (both Thermo Scientific). A compensation voltage of −40V and a total carrier gas flow of 3.5 L/min was used as well as an electrospray voltage of 1.9 kV was applied for ionization. The MS1 spectra was recorded using the Orbitrap analyzer at 120k resolution from m/z 380-980 using an automatic gain control (AGC) target of 500% and a maximum injection time of 3 ms. The Astral analyzer was used for MS/MS scans in data-independent mode with 2 Th non-overlapping isolation windows with a scan range of 150-2000 m/z. The precursor accumulation time was 3ms and an AGC target of 500%. The isolated ions were fragmented using HCD with 25% normalized collision energy.

### MS data acquisition of dimethylated peptides on the Orbitrap Astral

MS data acquisition was performed as described for mixed species samples on the Orbitrap Astral, if not described otherwise. For each of the six timepoints, triplicates of 50 ng of labeled peptide were injected. Samples were separated by the Whisper 40SPD method gradient (Evosep) on an Aurora Elite TS 15 cm and 75 µm ID (AUR3-15075C18-TS, IonOpticks) at 50 °C. The An electrospray voltage of 1.9 kV was applied. The MS1 resolution was 240 k with a maximum injection time of 100 ms and 6 ms for MS/MS.

### Data Analysis

All data analysis was performed with python 3.11 using Numpy, Pandas, Seaborn and Matplotlib.

### Search and analysis of dia-PASEF and synchro-PASEF data with alphaDIA

Data was searched with version 1.5.5 of alphaDIA using a previously published^32^ empirical HeLa library. A default single step search was used with the following parameters: *target_ms1_tolerance = 15ppm*, *target_ms2_tolerance = 15 ppm, target_candidates = 5*. For synchro-PASEF *quant_all = true* was set and a *quant_window* of 6 scans was used. All precursors with run-level FDR of 1% and protein groups with global FDR of 1% we’re accepted. Coefficients of variation we’re calculated on non-log transformed directLFQ normalized quantities.

### Search and analysis of ZenoTOF data with alphaDIA

Data was searched with version 1.5.5 of alphaDIA using the HeLa library mentioned above. A default single step search was used with the following parameters: *target_ms1_tolerance = 15ppm, target_ms2_tolerance = 15 ppm, target_candidates = 3, target_rt_tolerance = 300*. All precursors with run-level FDR of 1% and protein groups with global FDR of 1% we’re accepted. Coefficients of variation we’re calculated on non-log transformed directLFQ normalized quantities.

### Search and analysis of empirical library data from Lou et al

Raw files, libraries and fasta files were used as provided in the original publication^33^. All data was searched with alphaDIA 1.5.5 using default parameters. For timsTOF data the following parameters were changed: *target_ms1_tolerance = 15ppm, target_ms2_tolerance = 15 ppm, target_candidates = 5, quant_window = 6, group_level = genes, scans, target_rt_tolerance = 500* seconds. For QE-HF data search was performed with *target_ms1_tolerance = 5ppm, target_ms2_tolerance = 10 ppm, target_candidates = 5, quant_window = 6, group_level = genes, scans, target_rt_tolerance = 600* seconds. Data for benchmarked tools was used as provided in the original publication. Analysis was performed as described in the original publication except for reassignment of proteins. Instead, search engine specific protein grouping was used. For alphaDIA, precursor passing local 1% FDR and protein groups passing a global 1% FDR were accepted.

### Search and analysis of HeLa bulk data with fully predicted spectral libraries

For fully predicted library benchmarking, Spectronaut v18.6.231227.55695, DIA-NN 1.8.1, Chimerys on Ardia in Proteome discoverer and alphaDIA 1.5.4 was used. All analysis was performed using the same fasta file of reviewed human proteins without isoforms (01.12.2023). On all platforms, search was performed for tryptic precursors with carbamidomethyl modification at cysteine as fixed modification and variable methionine oxidation and protein N-terminal acetylation with maximum of two occurrences. Charge states 2 to 4 were included with sequence lengths between 7 and 35 amino acids with a single missed cleavage. For Chimerys, only peptides with up to 30 amino acids were used as the tool didn’t support 35 amino acids. For alphaDIA automatic library prediction by alphaPeptDeep was used using the Lumos model for a NCE of 25. AlphaDIA used default parameters for a two-step search with the following changes: *target_ms1_tolerance = 4 ppm, target_ms2_tolerance = 7 ppm, target_rt_tolerance = 300s* in the first pass and *target_rt_tolerance = 100s* for the second pass. All data was analyzed at a 1% FDR threshold as enforced by the search engine. Coefficients of variation we’re calculated on non-log intensities as provided by the search engine for all proteins. For Chimerys, quantification was only available on the protein level and not protein group level.

For Entrapment analysis, an Arabidopsis fasta with reviewed sequences and no isoforms was downloaded from Uniprot (02.02.2024). Search was performed as described above with heuristic inference. Following search all shared precursors, including isoleucine – leucine pairs were identified. Protein groups with shared precursors were discarded.

### Search and analysis of mixed species data with fully predicted spectral libraries

For all three species, reviewed non-isoform proteomes were downloaded from Uniprot (21.02.2024). Proteins were in-silico digested using tryptic cleavage with carbamidomethyl modification at cysteine as fixed modification and variable methionine oxidation and protein N-terminal acetylation with maximum of two occurrences. Charge states 2 to 4 were included with sequence lengths between 7 and 35 amino acids with a single missed cleavage. The Library was predicted using the alphaPeptDeep Lumos model at 25 NCE. AlphaDIA 1.5.4 was used with default parameters for a two-step search with the following changes: *target_candidates = 5, target_ms1_tolerance = 5 ppm, target_ms2_tolerance = 10 ppm, target_rt_tolerance = 200s* in the first pass and *target_rt_tolerance = 100s* for the second pass. Heuristic protein inference was used on the gene level. Proteins with shared sequences were removed as described above. For benchmarking accuracy, the median LFQ ratio was calculated for protein groups identified in at least three replicates.

### Search and analysis of SILAC data with fully predicted spectral libraries

A fully predicted human library was generated with alphaPeptDeep as described above but for a NCE of 27. The library was multiplexed across the light channel without additional modifications and a heavy channel with isotopic labeling of Arginine (+10.008269) and Lysine (+8.014199). A single step search was performed with alphaDIA default parameters apart from: *target_ms1_tolerance = 5ppm, target_ms2_tolerance = 20ppm, target_rt_tolerance = 600 seconds, channel_wise_fdr = True*.

### Search and analysis of dimethylated samples using transfer learning

A fully predicted human library was generated based on a reviewed human uniprot library (01.12.2023) with the general pretrained alphaPeptDeep model not trained on dimethylated peptides. The peptides were modified with Methionine oxidation and protein N-terminal acetylation as variable modifications with a maximum of two. N-Terminal and Lysine dimethylation were set as fixed modifications. Transfer search was performed using alphaDIA 1.5.5 with default parameters and *target_candidates = 1, target_ms1_tolerance = 4 ppm, target_ms2_tolerance = 7 ppm and target_rt_tolerance = 1200*. Transfer learning quantification was enabled and set to b and y ions with a maximum charge of 2 and the top 3 occurrences for every modified sequence. The generated transfer learning library was used for training with the default training scheme described above. For evaluation, the original pretrained model, the transfer learned retention time model, the transfer learned MS2 model and the fully transfer learned model were evaluated for search. All searches were performed with the same parameters as the transfer search apart from a *target_rt_tolerance = 100* for searches with the updated model.

### Search and analysis of transfer learning entrapments

For evaluation of transfer learning on FDRs, entrapment experiments with known false positive Arabidopsis peptides were performed on the unmodified HeLa bulk samples acquired on the Orbitrap Astral. The entrapment library was generated as described above for the two step search with added N-terminal glutamate and glutamine to pyroglutamate conversion as variable modification. Raw files were searched with alphaDIA 1.5.5 using default parameters and *target_candidates = 1, target_ms1_tolerance = 4 ppm, target_ms2_tolerance = 7 ppm and target_rt_tolerance = 1200*. Transfer learning quantification was enabled and set to b and y ions with a maximum charge of 2 and the top 3 occurrences for every modified sequence. Transfer learning was performed utilizing all human and Arabidopsis precursors identified at 1% FDR cutoff. The transfer learning model was then reused for a second search with updated *target_rt_tolerance = 150* seconds. The process was repeated twice and the identifications after every search were analyzed for the number of false positive Arabidopsis identifications as described above.

## Extended Data Figures

**Extended Data Fig. 1:**
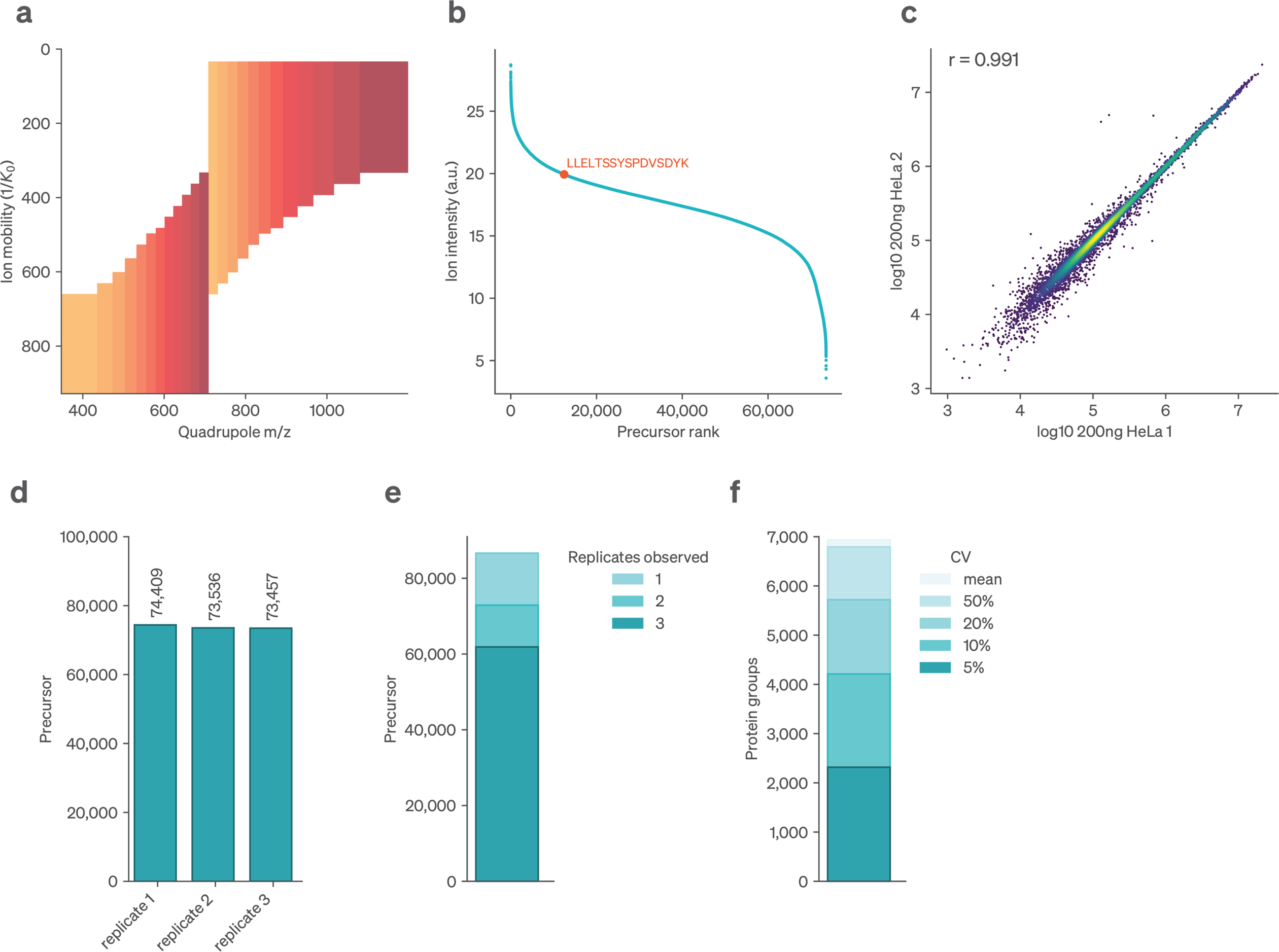
alphaDIA search results for library-based search of triplicate bulk HeLa dia-PASEF data. Data was acquired at 60SPD (21min) on the timsTOF Ultra. **a**, Overview of the MS2 window distribution scheme of optimal dia-PASEF. **b**, Precursor selected as example in Fig. 1 b-f. **c**, Correlation of LFQ protein quantities across replicates. **d**, number of precursors identified in each replicate at 1% FDR. **e**, Reproducibility of precursor identification across replicates. Number of precursors identified in at least 1, 2 or 3 replicates **f**, Precision of protein quantification. Number of protein groups for given CV cutoWs.

**Extended Data Fig. 2:**
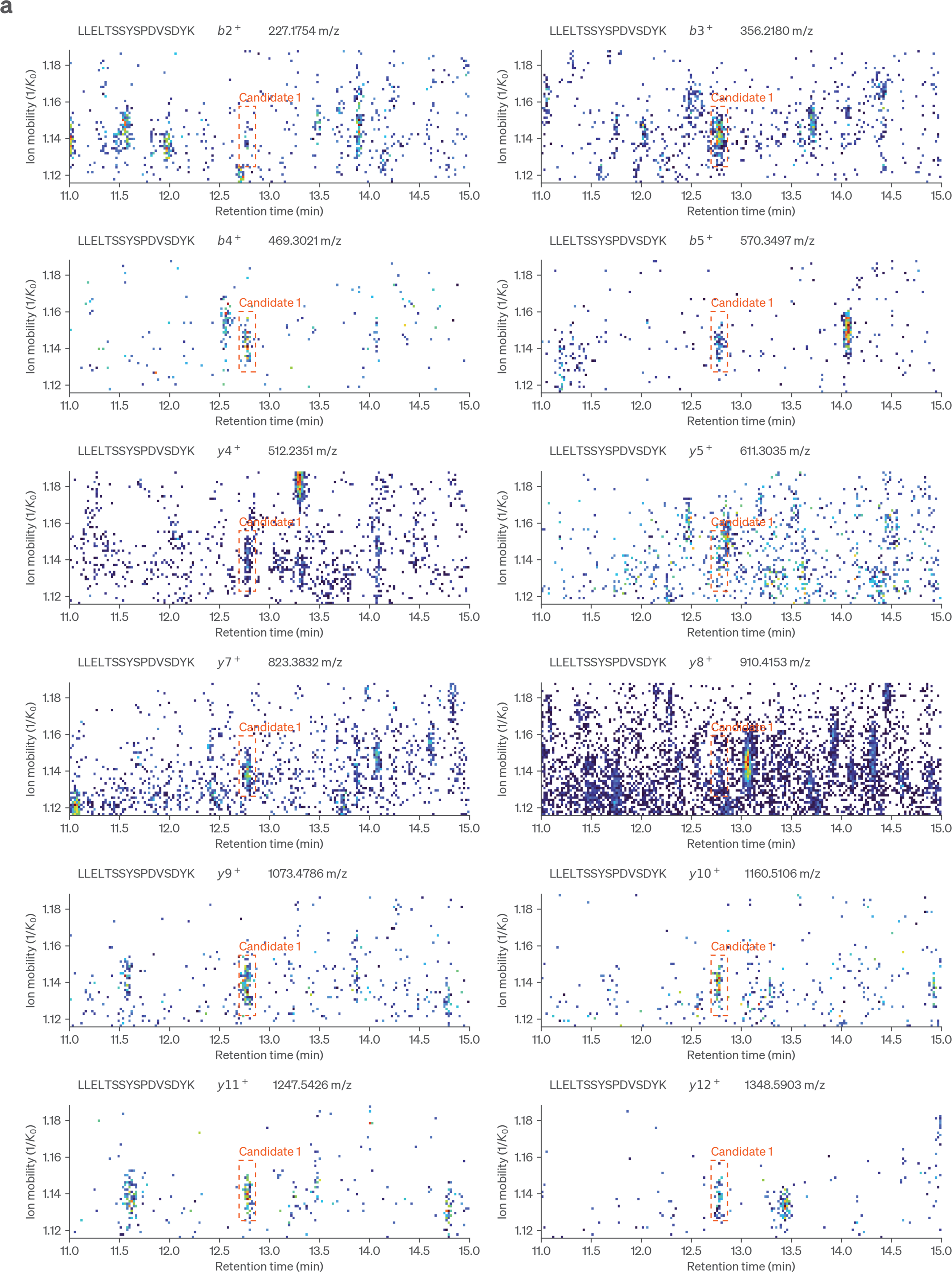
Fragment signal across ion mobility and retention time for the precursor LLELTSSYSPDVSDYK2+. **a**, For each fragment all signal within the 15ppm of calibrated mass tolerance is shown and the final integration boundaries of the identified precursor are highlighted in red. Due to the high sensitivity of time-of-flight detectors fragment signal might only correspond to few ion copies. This leads to stochastic sampling of ions and discontinuous signal across retention time and ion mobility. Distinguishing fragment signal from other ion species is challenging and prevents to determine clear peak boundaries. This requires an algorithm which does not need a minimum number of datapoints or certain peak shape. It’s likewise important to combine evidence across fragments for determination of peak group boundaries.

**Extended Data Fig. 3:**
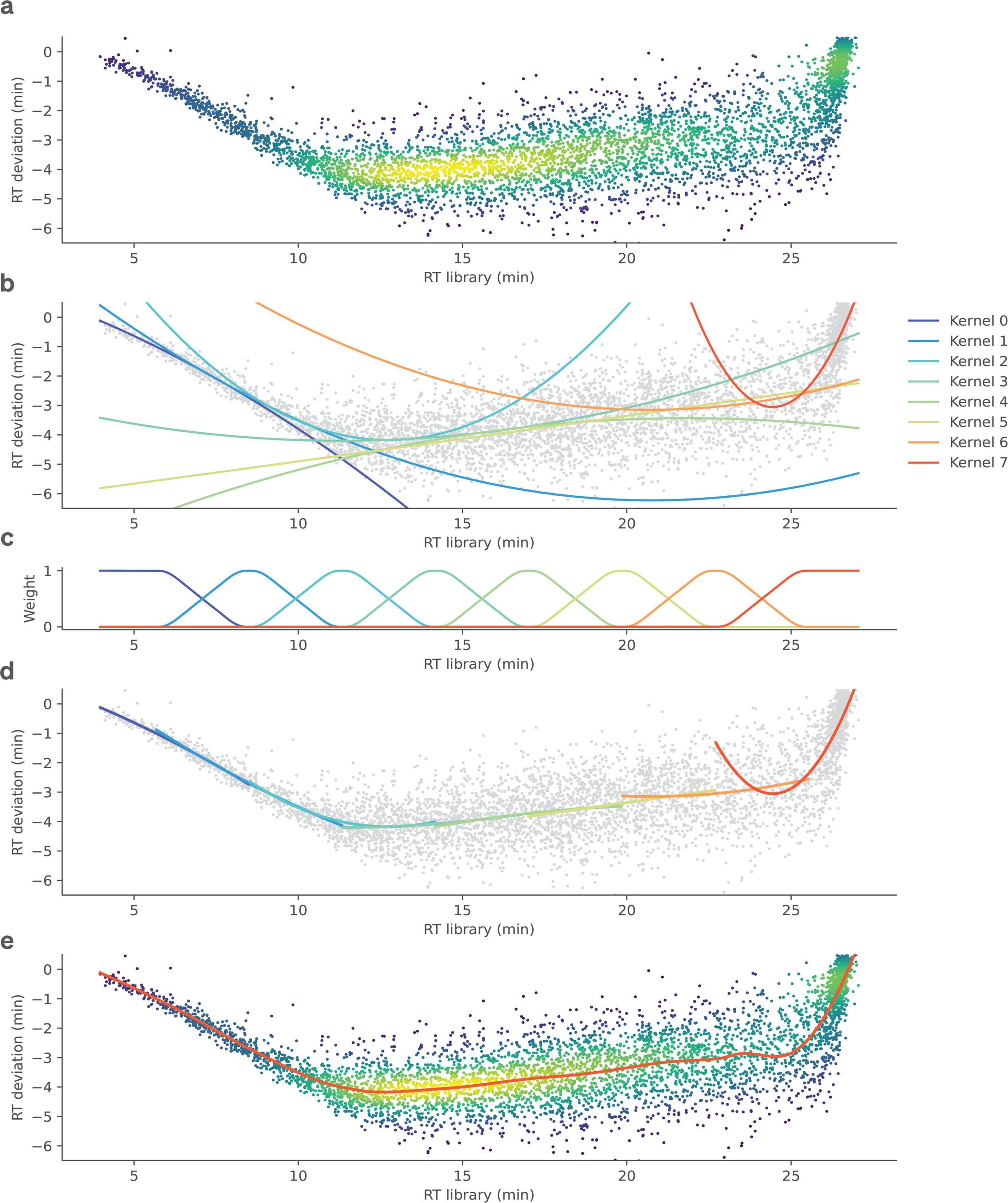
Calibration of library properties to observed data using locally estimated scatterplot smoothing (LOESS) regression. **a**, Observed retention times of confidently identified precursors compared with the library annotated values. The absolute deviation in minutes is shown. **b**, A collection of polynomial kernels is fitted to uniformly distributed subregions of the data. **c**, The functions are combined and smoothed using tricubic weights. **d**, Combining the kernels with their weighting functions allows to approximate the systematic deviation of the data locally. **e**, The sum of the weighted kernels can then be used for continuous approximation and calibration of retention times.

**Extended Data Fig. 4:**
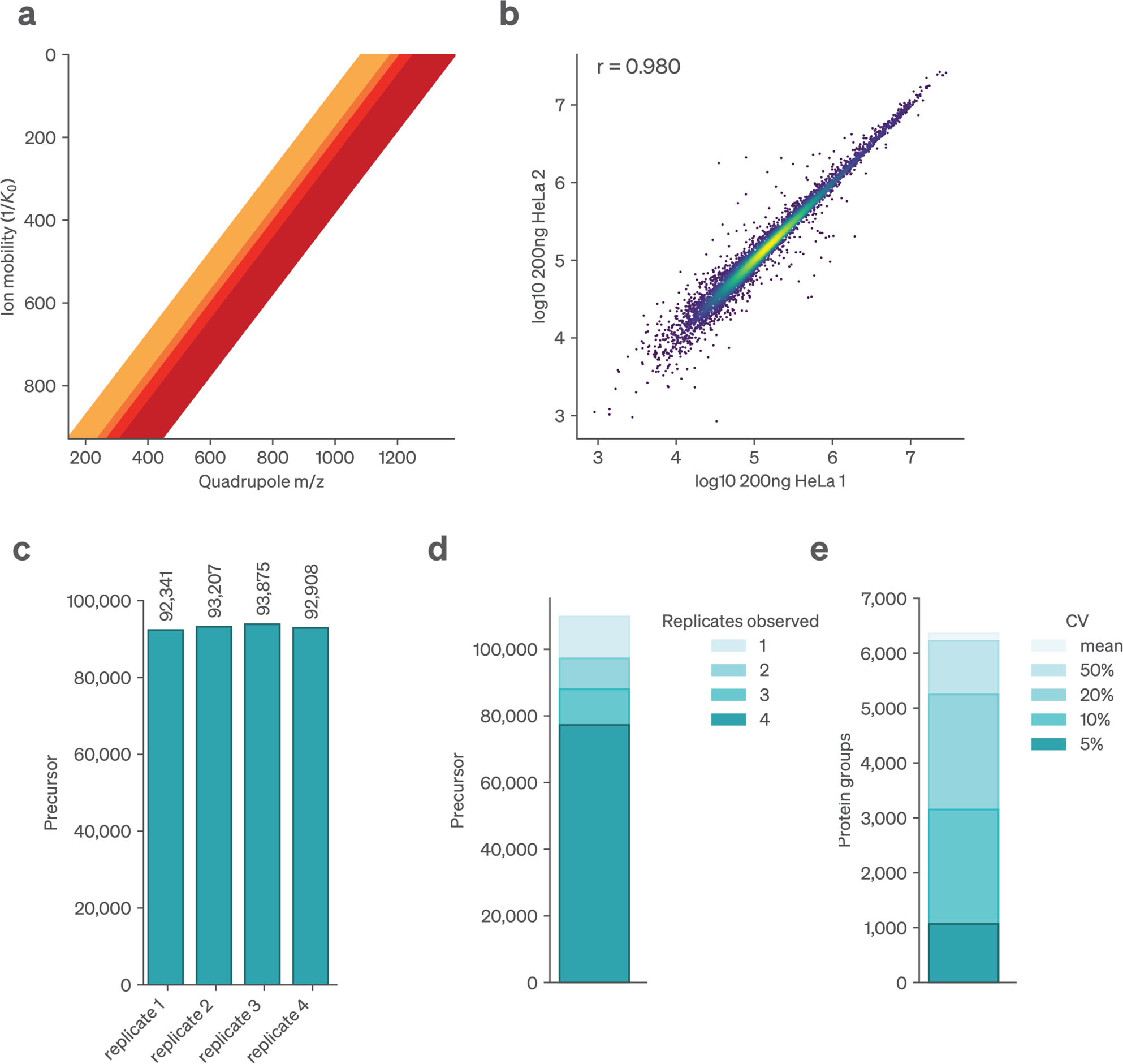
Processing of synchro-PASEF data with alphaDIA. Analysis of bulk HeLa lysate with synchro-PASEF on the timsTOF Ultra. **a,** In synchro-PASEF the quadrupole is continuously scanning across the mass range while ions elute from the TIMS trap. In this method, four synchro scans of variable width are being used. **b**, Correlation of protein groups quantified between two replicates of HeLa lysate **c**, Number of precursors identified at 1% FDR per replicate. **d**, Data completeness given by precursors identified in a minimum number of replicates. **e**, CoeWicient of variation (CV) for protein groups.

**Extended Data Fig. 5:**
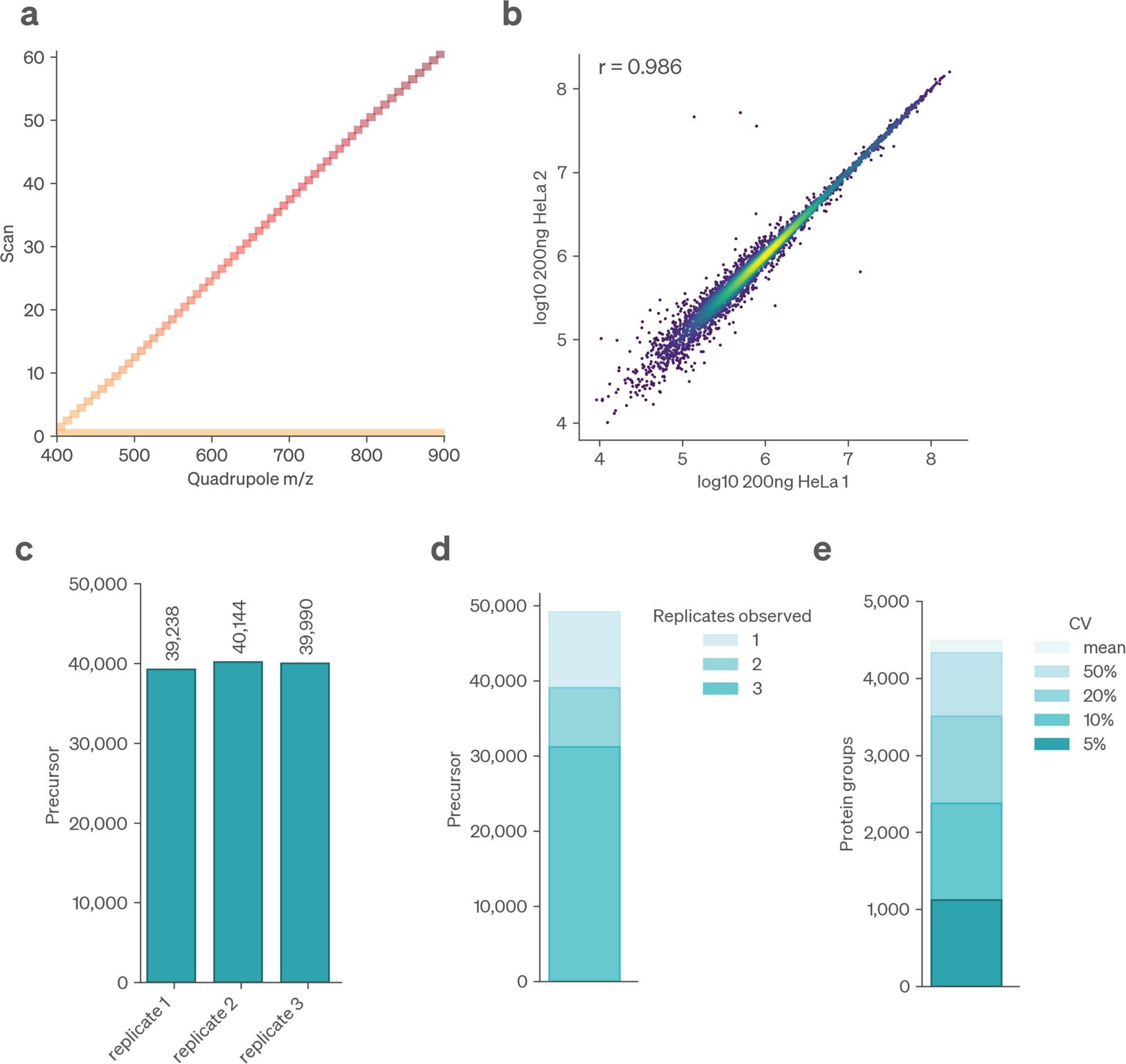
Analysis of Sciex swath data acquired on the ZenoTOF 7600. Bulk HeLa lysate was analyzed with 21minutes of active gradient. **a**, Overview of the acquisition method used for data acquisition. The position of MS2 quadrupole windows is shown for a single DIA cycle. **b**, Correlation of protein groups quantified between two replicates of HeLa lysate **c**, Number of precursors identified at 1% FDR per replicate. **d**, Data completeness given by precursors identified in a minimum number of replicates. **e**, CoeWicient of variation (CV) for protein groups.

**Extended Data Fig. 6:**
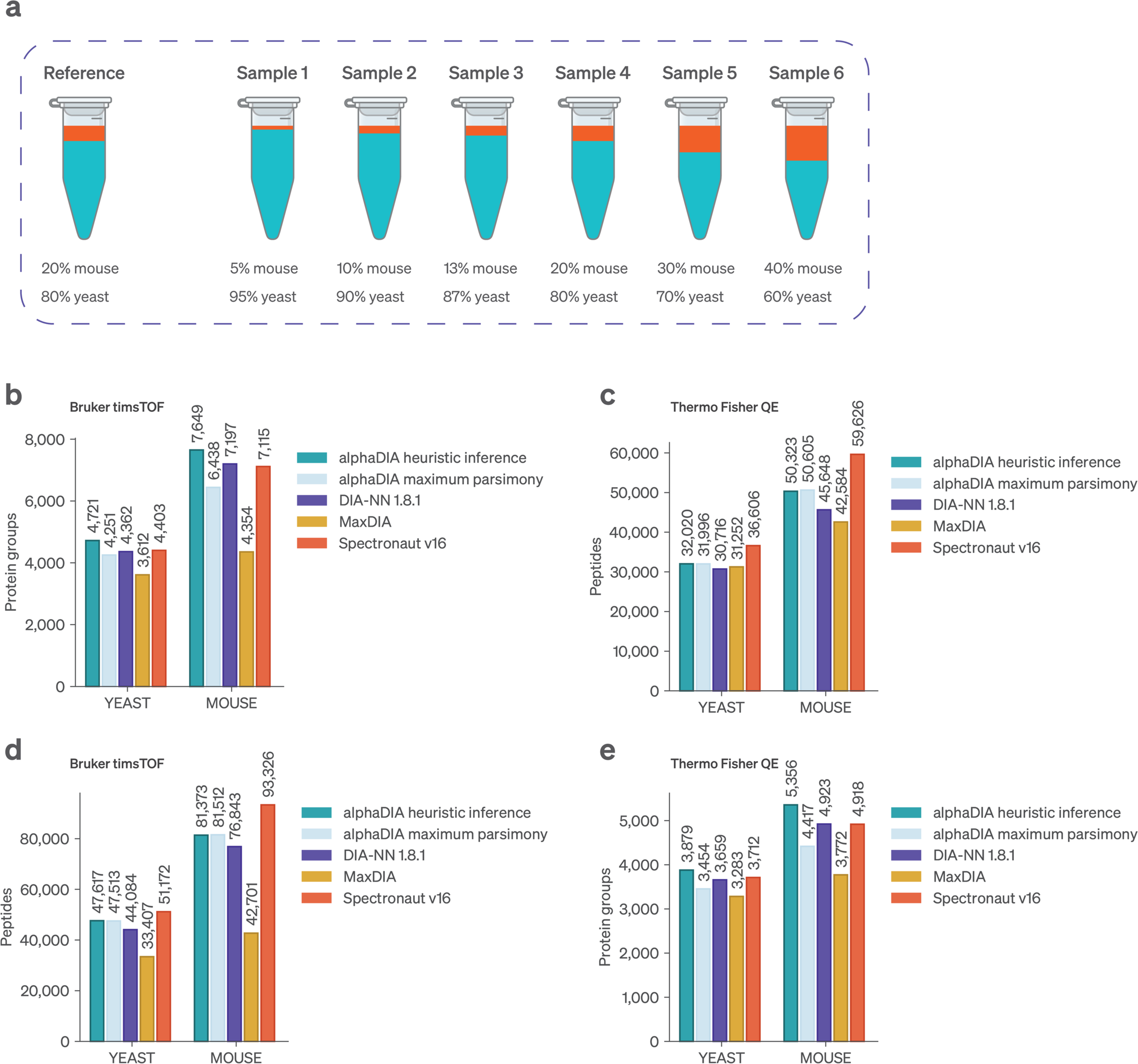
Benchmarking library based search in a complex background. **a**, Experimental setup as described by Lou et al.^33^ Mouse brain isolate digests were spiked into a complex yeast proteome background in diWerent ratios. **b**, Protein groups identified at 1% FDR on the Bruker timsTOF. **c**, Protein groups identified at 1% FDR on the Thermo Fisher QE-HF. **d**, Unique modified peptides identified 1% FDR across replicates on the Bruker timsTOF. **e**, Unique modified peptides identified 1% FDR across replicates on the Thermo Fisher QE-HF.

**Extended Data Fig. 7:**
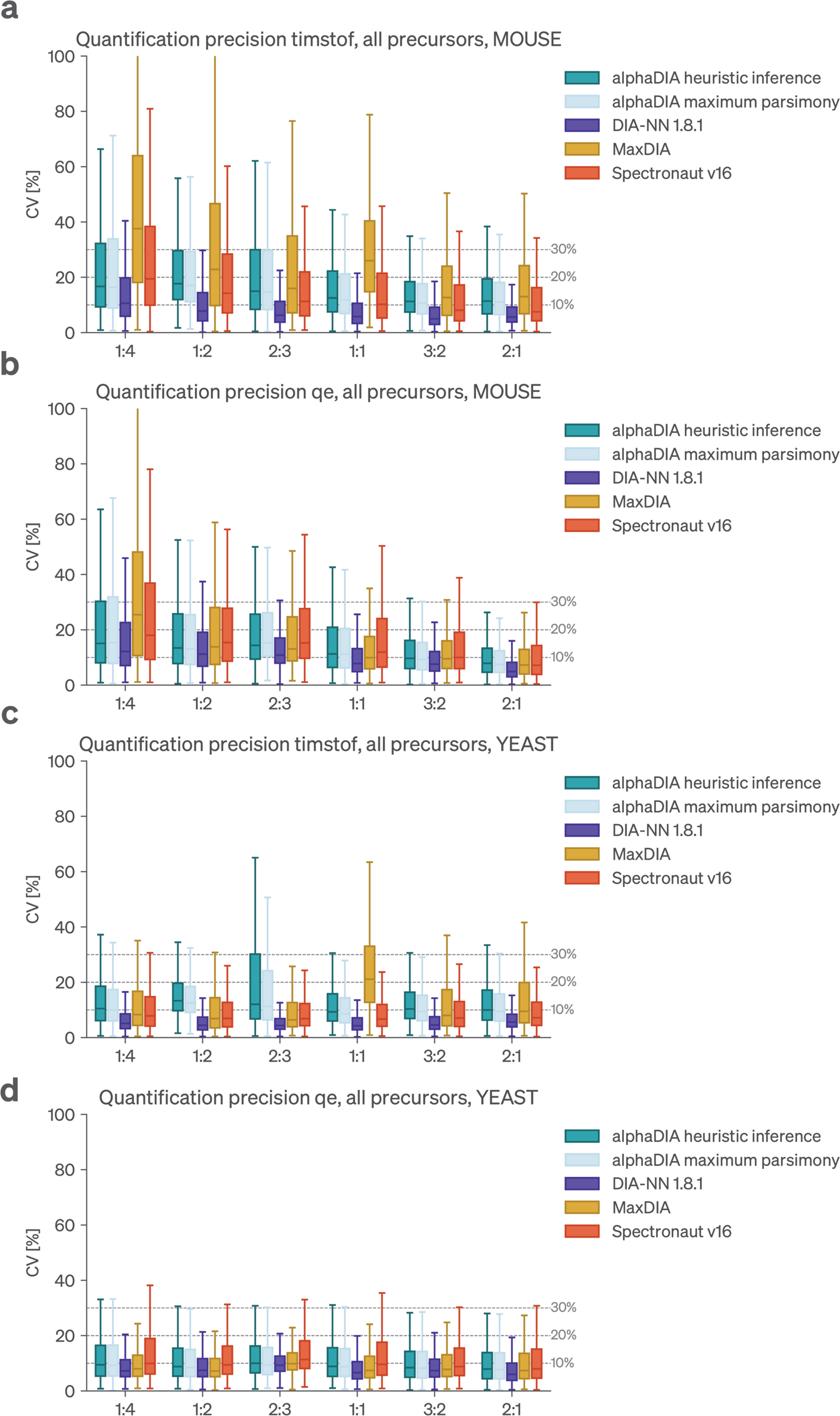
CoeJicient of variation for proteins in the empirical library benchmark. The quantitative precision was assessed by calculating the coeWicient of variation for quantifiable protein abundances, identified in at least three out of five replicates. **a**, Mouse proteins identified on the timsTOF. **b** Mouse proteins identified on the QE-HF **c**, Yeast proteins identified on the timsTOF. **d**, Yeast proteins identified on the QE-HF.

**Extended Data Fig. 8:**
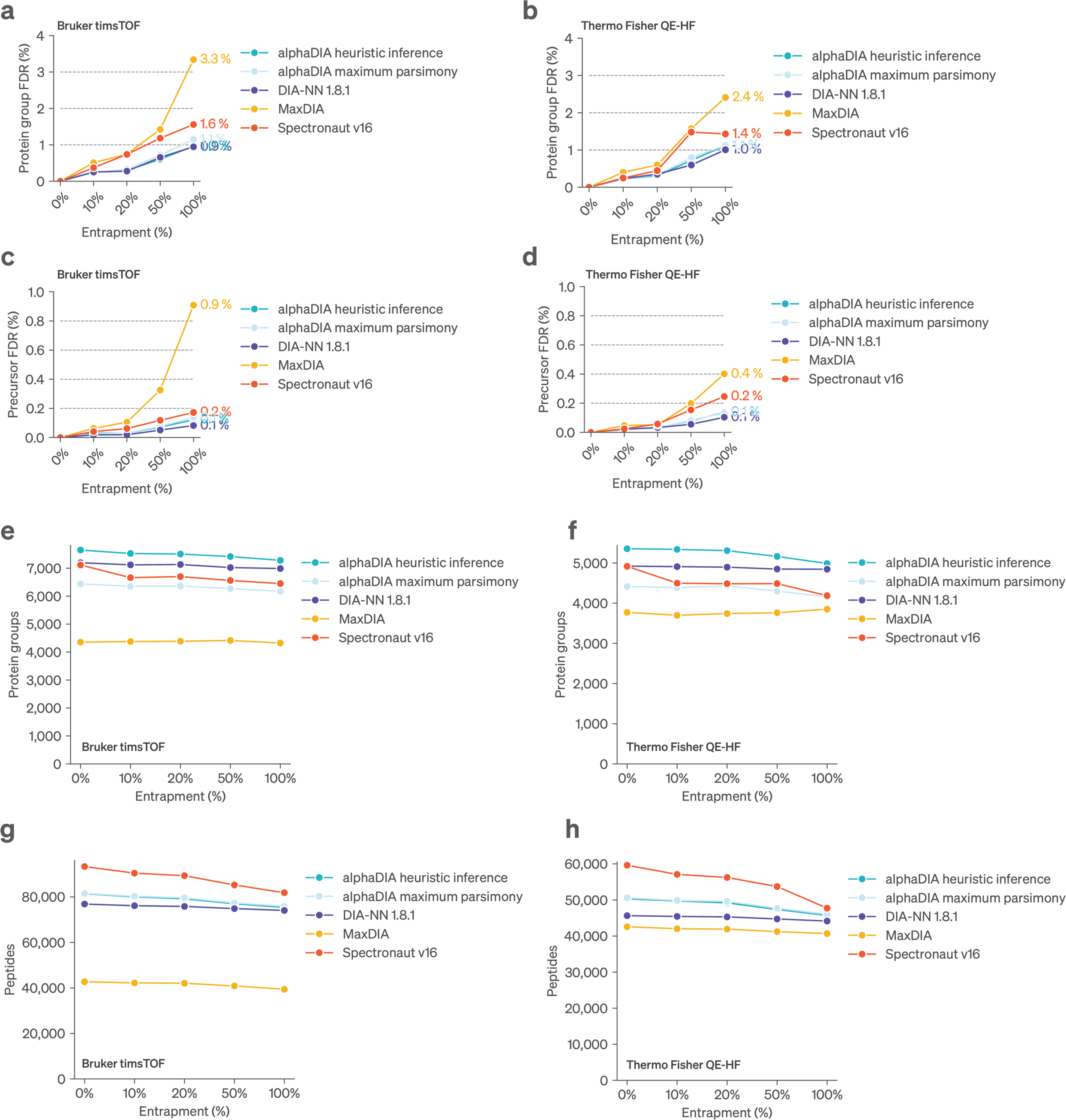
FDR benchmarking using Arabidopsis entrapments. Target Mouse and Yeast libraries we’re spiked in with increasing amounts of known false positive Arabidopsis precursors as provided by Lou et al.^33^ **a-d**, Number of global known false positive Arabidopsis proteins as a fraction of all identified proteins is shown as entrapment FDR. Search results are shown for increasing amounts of entrapment precursors, relative to the target library. **a**, Benchmarking data acquired on timsTOF, entrapment FDR calculated on the protein group level. **b**, Benchmarking data acquired on QE-HF, entrapment FDR calculated on the protein group level. **c**, Benchmarking data acquired on timsTOF, entrapment FDR calculated on the precursor level. **d**, Benchmarking data acquired on QE-HF, entrapment FDR calculated on the precursor level.

**Extended Data Fig. 9:**
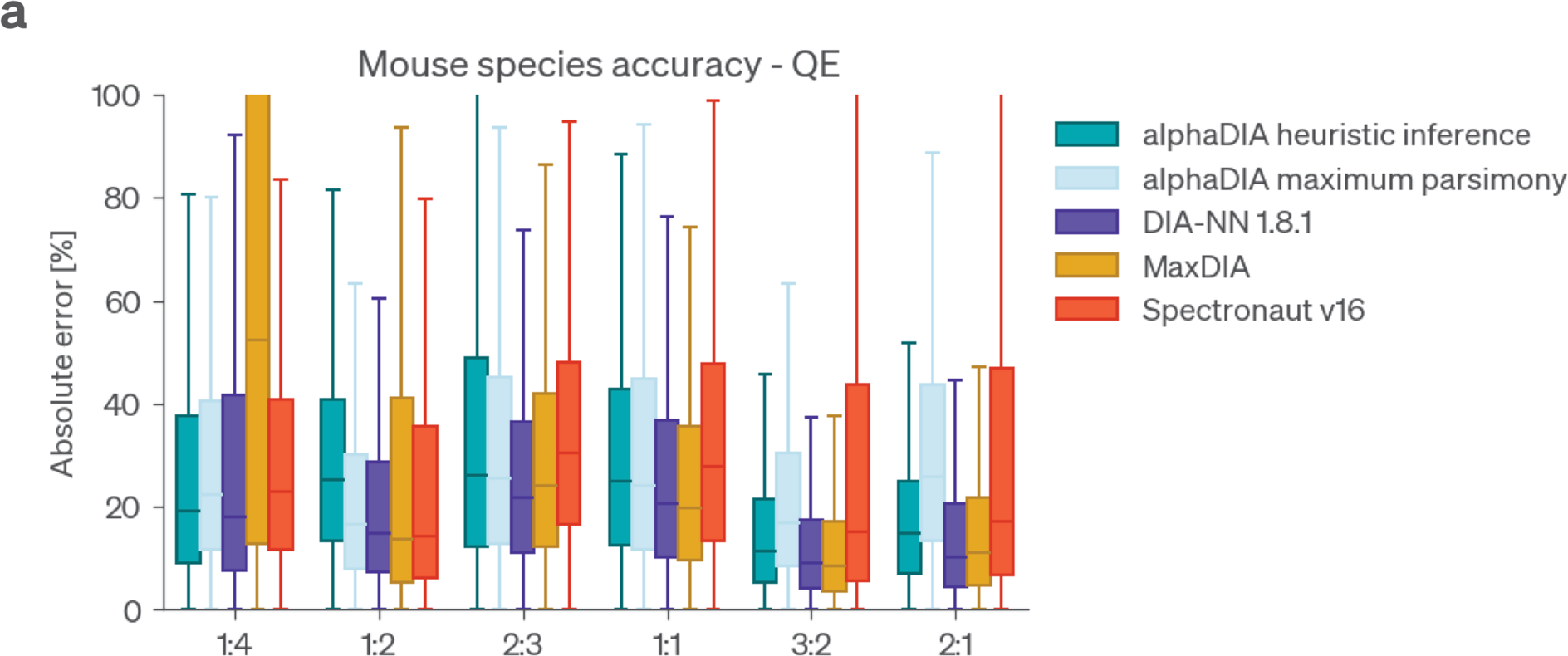
Quantitative accuracy for ratios in the benchmarking dataset. **a** Ratios were calculated as described in the original study. The absolute error between the expected and observed ratio is shown for diWerent search engines.

**Extended Data Fig. 10:**
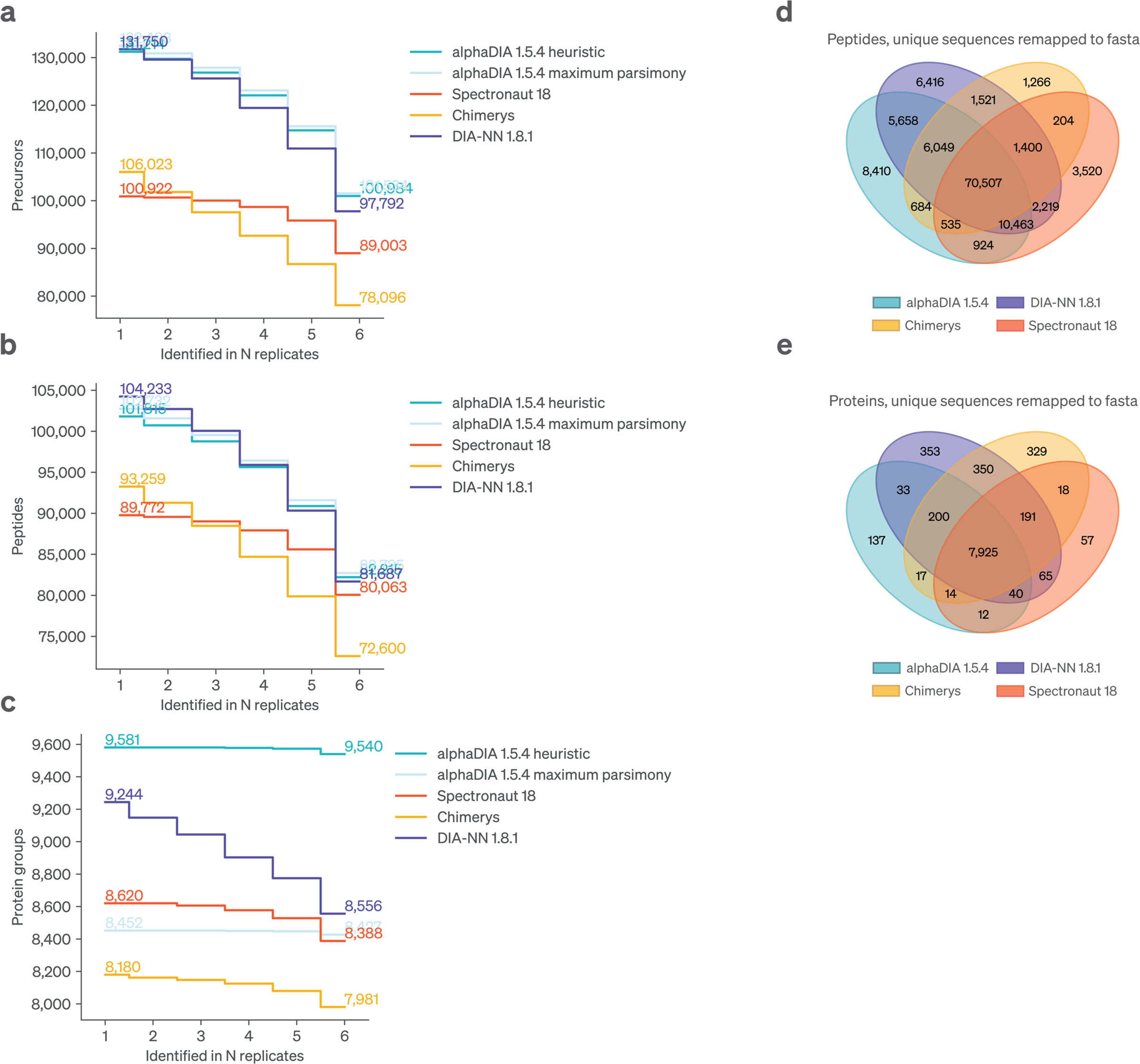
Comparison of identifications for fully predicted library search across search engines. **a**, Data completeness of precursor identifications across replicates. **b**, Data completeness of modified peptide identifications across replicates. **c**, Data completeness of protein identifications across runs. **d-e** Peptides were mapped back to the human reference proteome to enable comparison independent of grouping. All peptides matching to multiple proteins were discarded. **d**, Venn diagram comparing the peptides identified by the diWerent search engines. **e,** Venn diagram comparing the proteins identified by diWerent search engines.

**Extended Data Fig. 11:**
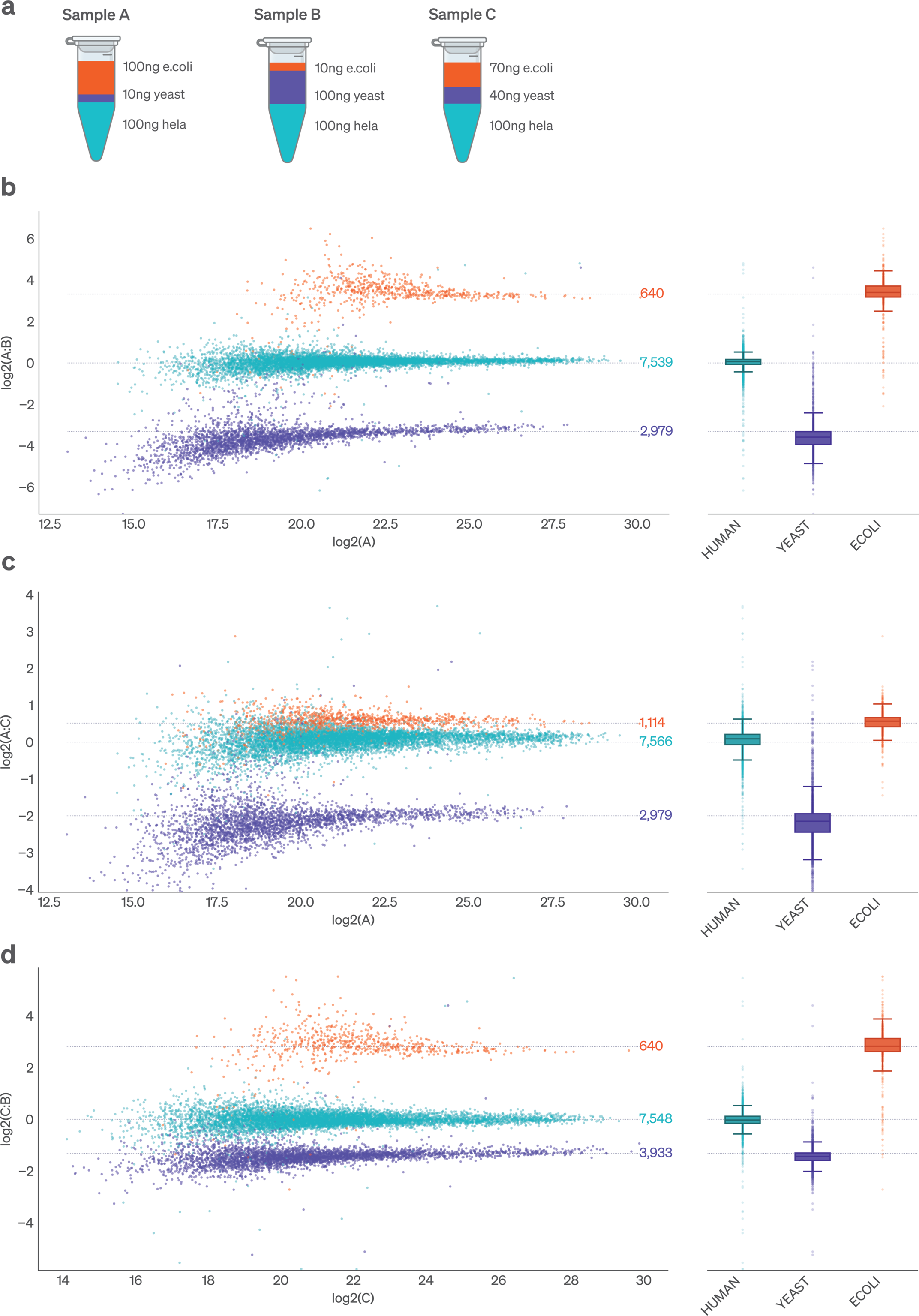
Quantitative accuracy benchmark using mixed species proteomes on the Orbitrap Astral. **a**, Five replicates of three samples were prepared with Yeast, E.coli and human proteomes mixed in defined ratios. **b**, Comparison of median protein group intensities at 1% FDR between sample A and B. **c**, Comparison of median protein group intensities at 1% FDR between sample A and B. **d**, Comparison of median protein group intensities at 1% FDR between sample C and B.

**Extended Data Fig. 12:**
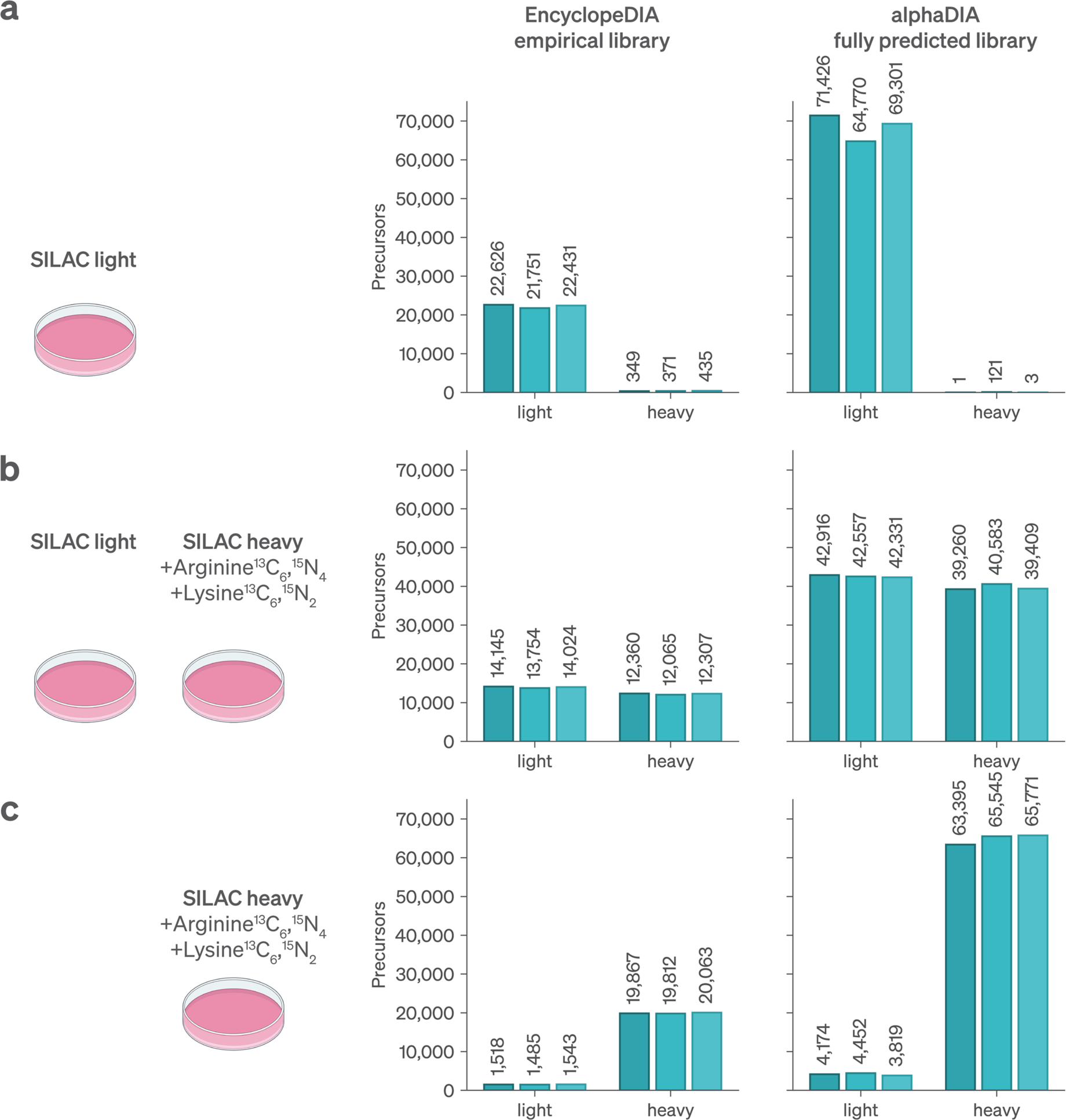
Validation of identification in SILAC labeled samples. SILAC data is from a method optimization study by the Garcia group that was originally analyzed by EncyclopeDIA and an empirical library^39^. This is compared to a fully alphaPeptDeep predicted library and database search by AlphaDIA. Triplicates results from the original paper are plotted in the left-hand panels and the AlphaDIA results on the same data in the right-hand panels. **a**, Percentage of false identifications in the heavy channel are median of 1.6% with EncyclopeDIA and 0.0043% with alphaDIA, which identified a threefold more precursors. **b**, For the combined sample, the heavy to light ratios are similar (46.7% heavy in EncyclopeDIA to 48.1% heavy in alphaDIA). **c**, After extended incorporation both analyses found similar percentage of light peptides (7.1% light in EncyclopeDIA vs 6.0% light in alphaDIA).

**Extended Data Fig. 13:**
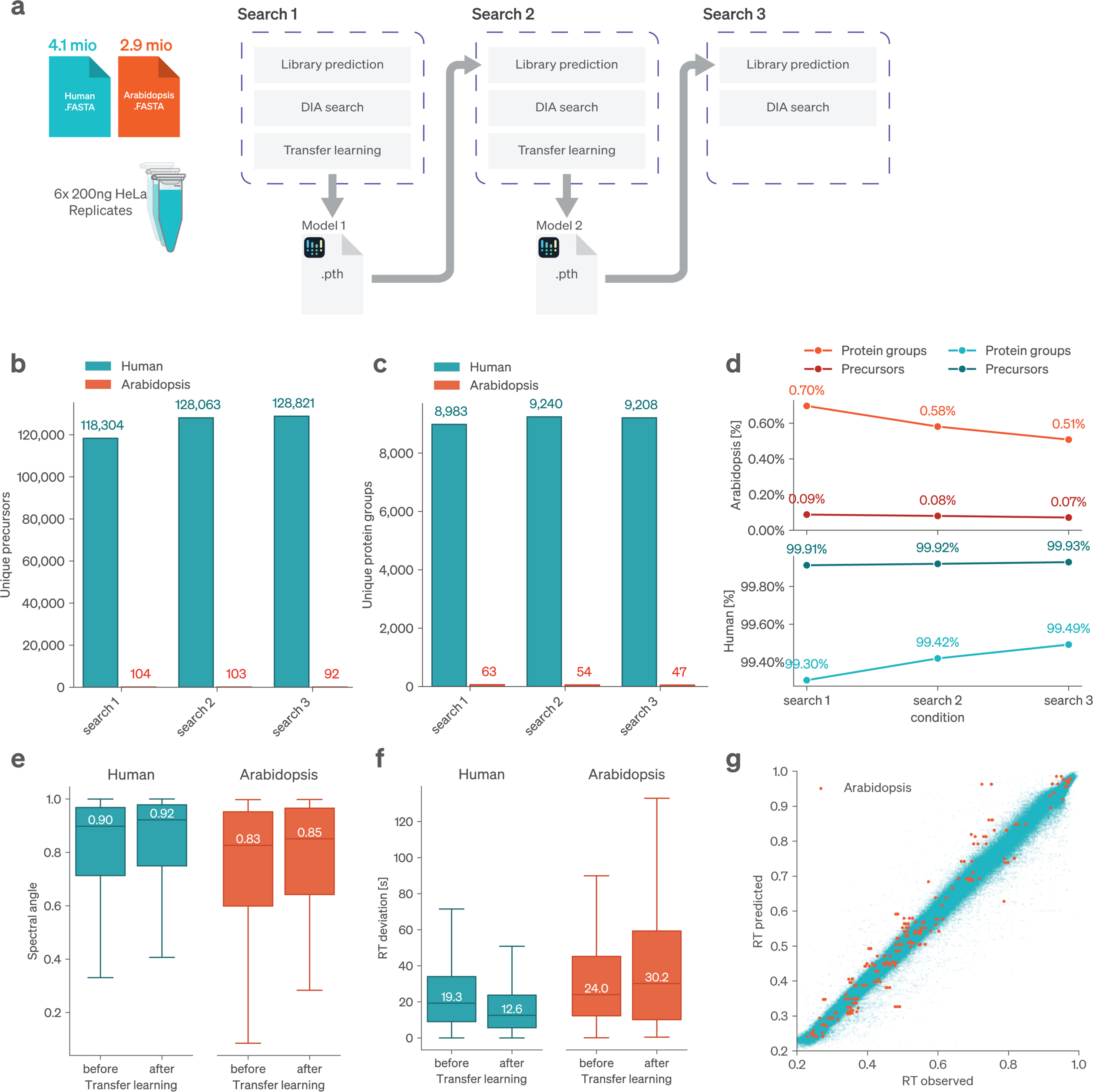
Entrapment validation of end-to-end transfer learning across for iterations. **a**, Overview of the validation workflow. A Human and Arabidopsis fasta file digest was used for fully predicted library search. All identified precursors at 1% FDR were subsequently used for end-to-end transfer learning, including false positive Arabidopsis identifications. This process was repeated twice, using the transfer learned deep-learning model for library prediction. **b**, Total unique identified precursors across six replicates. Precursors mapping to both species, including leucine and isoleucine pairs were removed. **c**, Total unique identified protein groups. **d**, Entrapment FDR given as the percentage of false positive Arabidopsis identifications. **e**, MS2 spectral angle for precursors before and after transfer learning. Median spectral angle is shown for each plot. **f**, Retention time deviation in seconds before and after transfer learning. The median retention time deviation is shown. **g**, Predicted vs observed retention time following transfer learning. False positive Arabidopsis identifications are highlighted in red.

**Extended Data Fig. 14:**
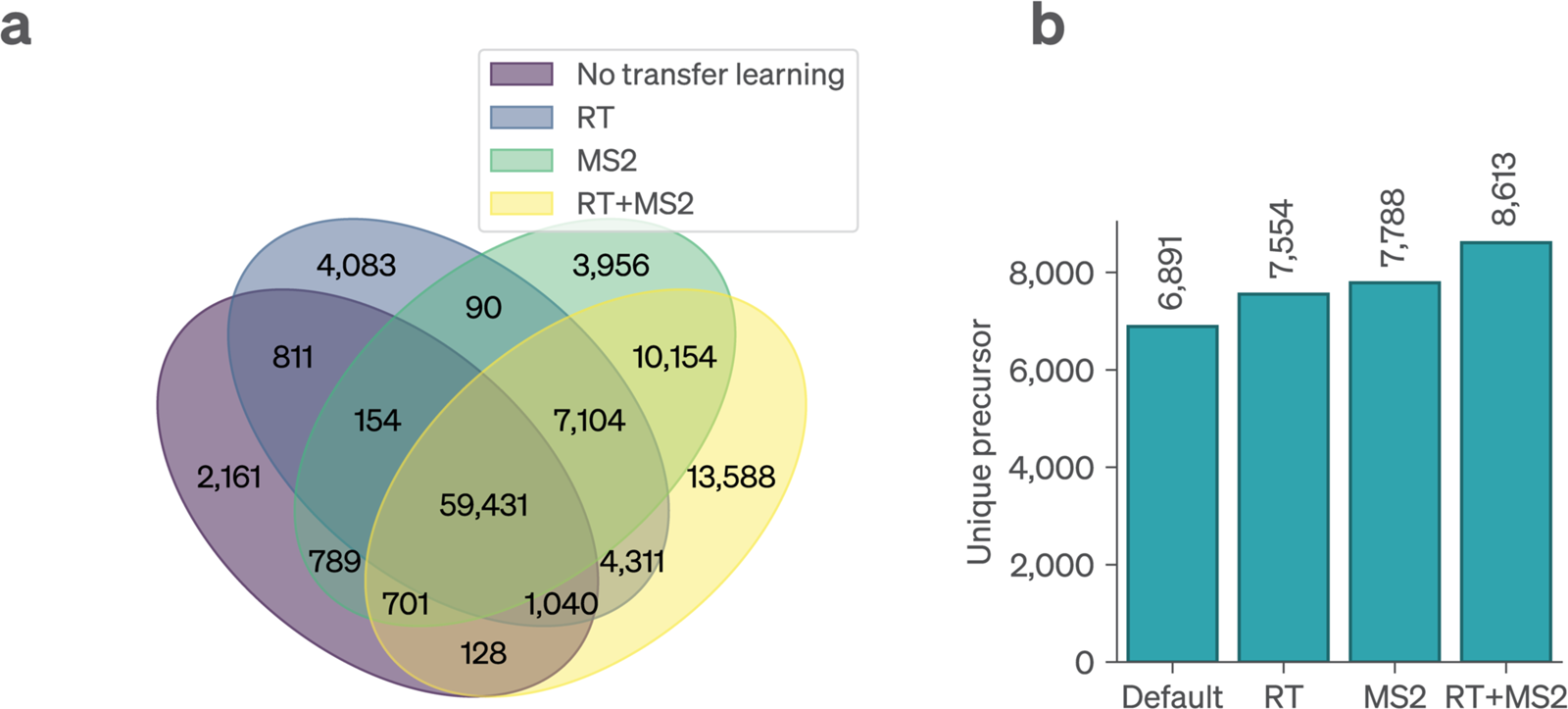
Comparison of identification with transfer learning of dimethylation. **a**, Venn diagram showing the overlap of precursor identifications before and after transfer learning. **b**, Total number of unique protein groups identified across replicates after diWerent stages of transfer learning.

## References

1. Aebersold, R. & Mann, M. Mass-spectrometric exploration of proteome structure and function. Nature 537, 347–355 (2016).

2. Navarro, P. et al. A multicenter study benchmarks software tools for label-free proteome quantification. Nat. Biotechnol. 34, 1130–1136 (2016).

3. Cox, J. & Mann, M. MaxQuant enables high peptide identification rates, individualized p.p.b.-range mass accuracies and proteome-wide protein quantification. Nat. Biotechnol. 26, 1367–1372 (2008).

4. Kong, A. T., Leprevost, F. V., Avtonomov, D. M., Mellacheruvu, D. & Nesvizhskii, A. I. MSFragger: ultrafast and comprehensive peptide identification in mass spectrometry–based proteomics. Nat. Methods 14, 513–520 (2017).

5. Lazear, M. R. Sage: An Open-Source Tool for Fast Proteomics Searching and Quantification at Scale. J. Proteome Res. 22, 3652–3659 (2023).

6. O’Reilly, F. J. & Rappsilber, J. Cross-linking mass spectrometry: methods and applications in structural, molecular and systems biology. Nat. Struct. Mol. Biol. 25, 1000–1008 (2018).

7. Virág, D. et al. Current Trends in the Analysis of Post-translational Modifications. Chromatographia 83, 1– 10 (2020).

8. Liu, H., Sadygov, R. G. & Yates, J. R. A Model for Random Sampling and Estimation of Relative Protein Abundance in Shotgun Proteomics. Anal. Chem. 76, 4193–4201 (2004).

9. Gillet, L. C. et al. Targeted Data Extraction of the MS/MS Spectra Generated by Data-independent Acquisition: A New Concept for Consistent and Accurate Proteome Analysis. Mol. Cell. Proteomics 11, O111.016717 (2012).

10. Collins, B. C. et al. Multi-laboratory assessment of reproducibility, qualitative and quantitative performance of SWATH-mass spectrometry. Nat. Commun. 8, 291 (2017).

11. Messner, C. B. et al. Ultra-fast proteomics with Scanning SWATH. Nat. Biotechnol. 39, 846–854 (2021).

12. Brunner, A. et al. Ultra-high sensitivity mass spectrometry quantifies single-cell proteome changes upon perturbation. Mol. Syst. Biol. 18, e10798 (2022).

13. Bernhardt, O. et al. Spectronaut: a fast and efficient algorithm for MRM-like processing of data independent acquisition (SWATH-MS) data. in (2014).

14. Tsou, C.-C. et al. DIA-Umpire: comprehensive computational framework for data-independent acquisition proteomics. Nat. Methods 12, 258–264 (2015).

15. Demichev, V., Messner, C. B., Vernardis, S. I., Lilley, K. S. & Ralser, M. DIA-NN: neural networks and interference correction enable deep proteome coverage in high throughput. Nat. Methods 17, 41–44 (2020).

16. Searle, B. C. et al. Chromatogram libraries improve peptide detection and quantification by data independent acquisition mass spectrometry. Nat. Commun. 9, 5128 (2018).

17. Sinitcyn, P. et al. MaxDIA enables library-based and library-free data-independent acquisition proteomics. Nat. Biotechnol. 39, 1563–1573 (2021).

18. Cox, J. Prediction of peptide mass spectral libraries with machine learning. Nat. Biotechnol. 41, 33– 43 (2023).

19. Gessulat, S. et al. Prosit: proteome-wide prediction of peptide tandem mass spectra by deep learning. Nat. Methods 16, 509–518 (2019).

20. Zeng, W.-F. et al. AlphaPeptDeep: a modular deep learning framework to predict peptide properties for proteomics. Nat. Commun. 13, 7238 (2022).

21. Bekker-Jensen, D. B. et al. Rapid and site-specific deep phosphoproteome profiling by data-independent acquisition without the need for spectral libraries. Nat. Commun. 11, 787 (2020).

22. Steger, M. et al. Time-resolved in vivo ubiquitinome profiling by DIA-MS reveals USP7 targets on a proteome-wide scale. Nat. Commun. 12, 5399 (2021).

23. Guzman, U. H. et al. Ultra-fast label-free quantification and comprehensive proteome coverage with narrow-window data-independent acquisition. Nat. Biotechnol. (2024) doi:10.1038/s41587-023-02099-7.

24. Wang, Z. et al. High-throughput proteomics of nanogram-scale samples with Zeno SWATH MS. eLife 11, e83947 (2022).

25. Meier, F. et al. diaPASEF: parallel accumulation–serial fragmentation combined with data-independent acquisition. Nat. Methods 17, 1229–1236 (2020).

26. Demichev, V. et al. dia-PASEF data analysis using FragPipe and DIA-NN for deep proteomics of low sample amounts. Nat. Commun. 13, 3944 (2022).

27. Distler, U. et al. midiaPASEF maximizes information content in data-independent acquisition proteomics. Preprint at 10.1101/2023.01.30.526204 (2023).

28. Skowronek, P. et al. Synchro-PASEF Allows Precursor-Specific Fragment Ion Extraction and Interference Removal in Data-Independent Acquisition. Mol. Cell. Proteomics 22, 100489 (2023).

29. Strauss, M. T. et al. AlphaPept: a modern and open framework for MS-based proteomics. Nat. Commun. 15, 2168 (2024).

30. Willems, S., Voytik, E., Skowronek, P., Strauss, M. T. & Mann, M. AlphaTims: Indexing Trapped Ion Mobility Spectrometry–TOF Data for Fast and Easy Accession and Visualization. Mol. Cell. Proteomics 20, 100149 (2021).

31. Ammar, C., Schessner, J. P., Willems, S., Michaelis, A. C. & Mann, M. Accurate Label-Free Quantification by directLFQ to Compare Unlimited Numbers of Proteomes. Mol. Cell. Proteomics 22, 100581 (2023).

32. Skowronek, P. et al. Rapid and In-Depth Coverage of the (Phospho-)Proteome With Deep Libraries and Optimal Window Design for dia-PASEF. Mol. Cell. Proteomics 21, 100279 (2022).

33. Lou, R. et al. Benchmarking commonly used software suites and analysis workflows for DIA proteomics and phosphoproteomics. Nat. Commun. 14, 94 (2023).

34. Huang, T., Wang, J., Yu, W. & He, Z. Protein inference: a review. Brief. Bioinform. 13, 586–614 (2012).

35. Granholm, V., Noble, W. S. & Käll, L. On Using Samples of Known Protein Content to Assess the Statistical Calibration of Scores Assigned to Peptide-Spectrum Matches in Shotgun Proteomics. J. Proteome Res. 10, 2671–2678 (2011).

36. Cox, J., et al. Accurate Proteome-wide Label-free Quantification by Delayed Normalization and Maximal Peptide Ratio Extraction, Termed MaxLFQ. Mol. Cell. Proteomics 13, 2513–2526 (2014).

37. Derks, J., et al. Increasing the throughput of sensitive proteomics by plexDIA. Nat. Biotechnol. 41, 50–59 (2023).

38. Thielert, M., et al. Robust dimethyl-based multiplex-DIA doubles single-cell proteome depth via a reference channel. Mol. Syst. Biol. 19, e11503 (2023).

39. Pino, L. K., Baeza, J., Lauman, R., Schilling, B. & Garcia, B. A. Improved SILAC Quantification with Data-Independent Acquisition to Investigate Bortezomib-Induced Protein Degradation. J. Proteome Res. 20, 1918–1927 (2021).

40. Nesvizhskii, A. I. A survey of computational methods and error rate estimation procedures for peptide and protein identification in shotgun proteomics. J. Proteomics 73, 2092–2123 (2010).

41. Nesvizhskii, A. I. & Aebersold, R. Interpretation of Shotgun Proteomic Data. Mol. Cell. Proteomics 4, 1419–1440 (2005).

